# Population receptive field estimates for motion-defined stimuli

**DOI:** 10.1101/435735

**Authors:** Anna E. Hughes, John A. Greenwood, Nonie J. Finlayson, D. Samuel Schwarzkopf

## Abstract

The processing of motion changes throughout the visual hierarchy, from spatially restricted ‘local motion’ in early visual cortex to more complex large-field ‘global motion’ at later stages. Here we used functional magnetic resonance imaging (fMRI) to examine spatially selective responses in these areas related to the processing of random-dot stimuli defined by differences in motion. We used population receptive field (pRF) analyses to map retinotopic cortex using bar stimuli comprising coherently moving dots. In the first experiment, we used three separate background conditions: no background dots (dot-defined bar-only), dots moving coherently in the opposite direction to the bar (kinetic boundary) and dots moving incoherently in random directions (global motion). Clear retinotopic maps were obtained for the bar-only and kinetic-boundary conditions across visual areas V1-V3 and in higher dorsal areas. For the global-motion condition, retinotopic maps were much weaker in early areas and became clear only in higher areas, consistent with the emergence of global-motion processing throughout the visual hierarchy. However, in a second experiment we demonstrate that this pattern is not specific to motion-defined stimuli, with very similar results for a transparent-motion stimulus and a bar defined by a static low-level property (dot size) that should have driven responses particularly in V1. We further exclude explanations based on stimulus visibility by demonstrating that the observed differences in pRF properties do not follow the ability of observers to localise or attend to these bar elements. Rather, our findings indicate that dorsal extrastriate retinotopic maps may primarily be determined by the visibility of the neural responses to the bar relative to the background response (i.e. neural signal-to-noise ratios) and suggests that claims about stimulus selectivity from pRF experiments must be interpreted with caution.

## 1. Introduction

Motion perception is one of the fundamental dimensions of vision **(Nakayama, 1985; Nishida, 2011)**, and it is now known that many areas of the brain are involved in motion processing **(Dupont et al., 1994; Pitzalis et al., 2010; Sunaert et al., 1999; Tootell et al., 1997)**. Converging psychophysical, electrophysiological, and imaging evidence suggests that motion is processed in an hierarchical manner, with signals first being processed locally (within restricted spatial windows) in areas such as V1 and then combined at higher levels in the visual cortical hierarchy to generate global motion percepts over larger regions of the visual field **(Adelson and Movshon, 1982; Braddick et al., 2001; Van Essen and Gallant, 1994; Williams and Sekuler, 1984)**.

Many psychophysical studies have used tasks involving the detection or discrimination of coherent motion to study the distinction between local-and global-motion processing **(Britten et al., 1993, 1992; Newsome and Paré, 1988; Scase et al., 1996; Watamaniuk, 1993)**. Due to the aperture problem, direction-selective neurons in lower visual areas such as V1 are thought to process only local motion – the 1D motion orthogonal to the orientation of the edge that is passing through the receptive field **(Adelson and Movshon, 1982; Marr and Ullman, 1981; Movshon, 1986; Wallach, 1935)**. To be able to process global motion, these 1D signals must be integrated over a relatively wide visual field area/region, a process thought to occur in higher visual areas, such as V5/MT+ to generate a global motion direction **(Heeger et al., 1996; Simoncelli and Heeger, 1998)**. fMRI evidence has provided support for a distinction in the neural locations of local and global processing, showing that V1 was activated more by incoherent local noise than coherent global motion, perhaps because the noise stimulus led to the activation of neurons with a wider range of motion selectivities. The reverse pattern was seen in V5 and V3A, which both responded more to coherent motion compared to the noise stimulus **(Braddick et al., 2001)**.

Other brain regions may be specialized for the detection of more complex motion patterns. One example is that of kinetic boundaries, where an edge is defined by differences in coherent motion direction on either side of the edge. A specific brain region known as the Kinetic Occipital area (KO), which is thought to include areas such as V3, V3A and V3B **(Larsson and Heeger, 2006)**, may be specialized for detecting these boundaries **(Van Oostende et al., 1997)**. However, other research suggests that KO may not be completely specialized for motion-boundary processing, with KO thought to also respond preferentially to stimuli such as form cues **(Zeki et al., 2003)**. It has further been shown that other visual areas such as LO1, LO2 and V7 respond preferentially to motion boundaries, suggesting that motion-boundary processing may be more widely distributed across the visual cortex **(Larsson et al., 2010)**.

Despite this debate over the specificity of the area responding to kinetic contours, it is clear that kinetic boundaries are relatively complex stimuli that are not processed at lower levels in the visual hierarchy: they produce little fMRI response in lower visual areas such as V1 and V2 **(Van Oostende et al., 1997; Zeki et al., 2003)**, consistent with electrophysiological evidence that the majority of neurons in these areas do not respond to kinetic contours **(Leventhal et al., 1998; Marcar et al., 2000)**.

The visual hierarchy also has multiple representations of the retina, laid out in topological maps that are commonly called retinotopic maps **(Felleman and Van Essen, 1991; Sereno et al., 1995)**. While it was once thought that retinotopy was a property of lower level visual areas, it is now known that areas such as MT+ and MST also contain retinotopic maps (known as TO1 and TO2 respectively; **(Amano et al., 2009)**, as do the frontal eye fields **(Kastner et al., 2007)** and even the default mode network **(Knapen et al., 2018)**. Indeed, despite the large receptive field sizes within primate MT/V5, it is possible to track object position at the population level **(Chen et al., 2015)**. This suggests that retinotopy is a general organizing principle within the cortex. It is therefore of interest to know whether retinotopic map properties vary according to the visual area under question, and particularly whether these properties are affected by the different functional selectivities of different regions of the brain.

One technique that has been used to analyse retinotopic maps via fMRI is population receptive field (pRF) analysis, providing an estimation of both the visual field position preferred by each voxel and the range of visual field locations where a stimulus evokes a response **(Dumoulin and Wandell, 2008)**. pRFs can therefore be thought of as a statistical summary of the neuronal properties within a sampled region. Recent work has shown systematic differences in pRF sizes across different brain regions and eccentricities, with size increasing along the visual processing hierarchy and with increasing eccentricity **(Alvarez et al., 2015; Amano et al., 2009; Dumoulin and Wandell, 2008; Haas et al., 2014; Harvey and Dumoulin, 2011; Schwarzkopf et al., 2014)**.

Across these different brain areas, several factors can affect pRF size and position. Retinotopy in early visual areas is primarily thought to be stimulus-driven, but there is evidence that higher level maps can also be attentionally-driven **(Tootell et al., 1998)**. One study **(Saygin and Sereno, 2008)** used point light biological walkers moving in a ‘wedge’ stimulus to dissociate attention and stimulus effects in retinotopic mapping, and showed that V1 did not respond clearly when the distinction between stimulus and background was driven only attentionally, but it did respond when there was a visual difference between stimulus and background without attention directed to the stimulus. The opposite pattern was seen in frontal and parietal areas. The spatial tuning of pRFs is also affected by attentional load at fixation, with pRF size increasing and pRF location becoming more eccentric under high perceptual load **(Haas et al., 2014)**. Similar results have also been reported in face-selective brain regions **(Kay et al., 2015)**. This position modulation seems to occur across the entire visual field, not just at the attended location **(Klein et al., 2014)**, and recent work suggests that these position shifts are the key mechanism by which attention enhances discriminability and representational quality of stimuli **(Vo et al., 2017)**. It therefore seems that attention can affect cortical spatial tuning properties, in turn altering the visibility of stimulus differences.

Stimulus properties may also alter the measured properties of pRFs. In particular, the fMRI response of a voxel may be driven by different groups of cells depending upon the properties of the stimulus. While most studies have used simple luminance-defined stimuli (such as checkerboards) to generate pRF maps, more complex stimuli may be especially suited for generating maps in higher visual areas **(Yildirim et al., 2018)**. A recent study that used a pRF mapping stimulus designed specifically to isolate orientation contrast showed reductions in measured pRF size in higher visual areas such as LO compared to the measurements made with standard luminance-based stimuli **(Yildirim et al., 2018)**. pRF sizes have also been shown to vary based on the alignment and curvature of contours within mapping stimuli **(Dumoulin et al., 2014)**. In the motion domain, there is evidence that MT+ may be more susceptible to stimulus configuration than earlier visual areas **(Alvarez et al., 2015)** but to date there has been no systematic investigation of the effects of different motion stimuli on pRF measurement.

Given the above variations due to stimulus properties and attentional state, in this study we asked whether stimuli thought to preferentially drive distinct stages of the motion-processing hierarchy can similarly alter the estimation of pRF parameters. In our initial experiment, we tested this with a moving bar stimulus similar to those commonly used in pRF mapping studies, defined by dots moving coherently. We then used different backgrounds in an attempt to differentially drive responses in different brain regions. In the bar-only condition, the bar was presented alone against a grey background. We predicted that this stimulus would generate a strong visual signal and enable the generation of pRF maps at all levels of the motion-processing hierarchy, much like a typical pRF mapping stimulus. In the ‘kinetic’ condition the bar was defined by kinetic boundaries. We predicted that if these stimuli are preferentially processed in KO/V3B, we might expect smaller pRF sizes and a higher proportion of voxels responding in this area. Finally, in the ‘global’ condition, the background consisted of incoherently moving dots. We therefore predicted that for the ‘global’ condition, higher visual areas that process stimuli in a more global manner should be able to distinguish between the bar and background and thus generate good pRF maps. In contrast, for these latter two conditions, we predicted that V1 would not be able to distinguish the bar from the background, leading to a reduced response.

In a second experiment, we asked whether any differential responses seen in the first experiment were a consequence of differences in the selectivity of these motion-selective regions, or whether they could be explained by other factors, such as the visibility of the stimulus. We compared a bar-only stimulus to two conditions with lower visibility bars: one motion-defined stimulus, where the bar was defined by transparent motion (against a non-transparent background), and a non-motion defined stimulus, where the bar was instead defined by differences in dot size. If pRF properties varied due to differential motion processing, we predicted that a different pattern of responses should arise for these two stimuli: for example, the ‘transparent’ stimulus should show lower responsivity than the ‘size defined’ stimulus in V1, but higher responsivity in higher visual areas selective for global motion. In a similar vein, the subtle dot size difference in the size-defined stimulus should maximize the signal in V1 and the early visual cortex compared to higher regions. However, if the differential responses were simply due to the visibility of the bar stimulus, we predicted similar responses for the two conditions across different visual areas. We also conducted a third behavioural experiment as a control, presenting each of the above mapping stimuli and requiring observers to localise the bar element. This allowed us to further assess whether variations in the properties of pRFs could be predicted by the visibility of the bar stimuli, or whether these variations can be attributed to the visibility of the neural signals underlying the BOLD response.

## 2. Experiment 1

Here we examined retinotopic maps and pRF properties across the visual hierarchy using three distinct retinotopic mapping stimuli: the ‘bar-only’ stimulus, similar to standard retinotopic mapping stimuli; the ‘global’ bar stimulus with coherent motion against a background of noise; and the ‘kinetic’ bar stimulus with coherent motion against a background of oppositely-moving dots.

### 2.1. Materials and Methods

#### 2.1.1 Participants

Five participants (one male) took part in Experiment 1, including two of the authors (AH and JG) and three experienced participants <u>naïve</u> to the aims of the study. Participants were aged between 24-36 years (mean = 29.6 years) and one participant was left handed. All participants were experienced in an fMRI context and had normal or corrected-to-normal visual acuity. Written consent was acquired from all participants to ensure that they understood the potential risks associated with fMRI. The experiments were approved by the UCL Research Ethics Committee.

#### 2.1.2 Stimuli

Figure 1 shows a schematic of the experimental set up, and GIF versions of the different experimental stimuli are available as supplementary material.

**Figure 1.**
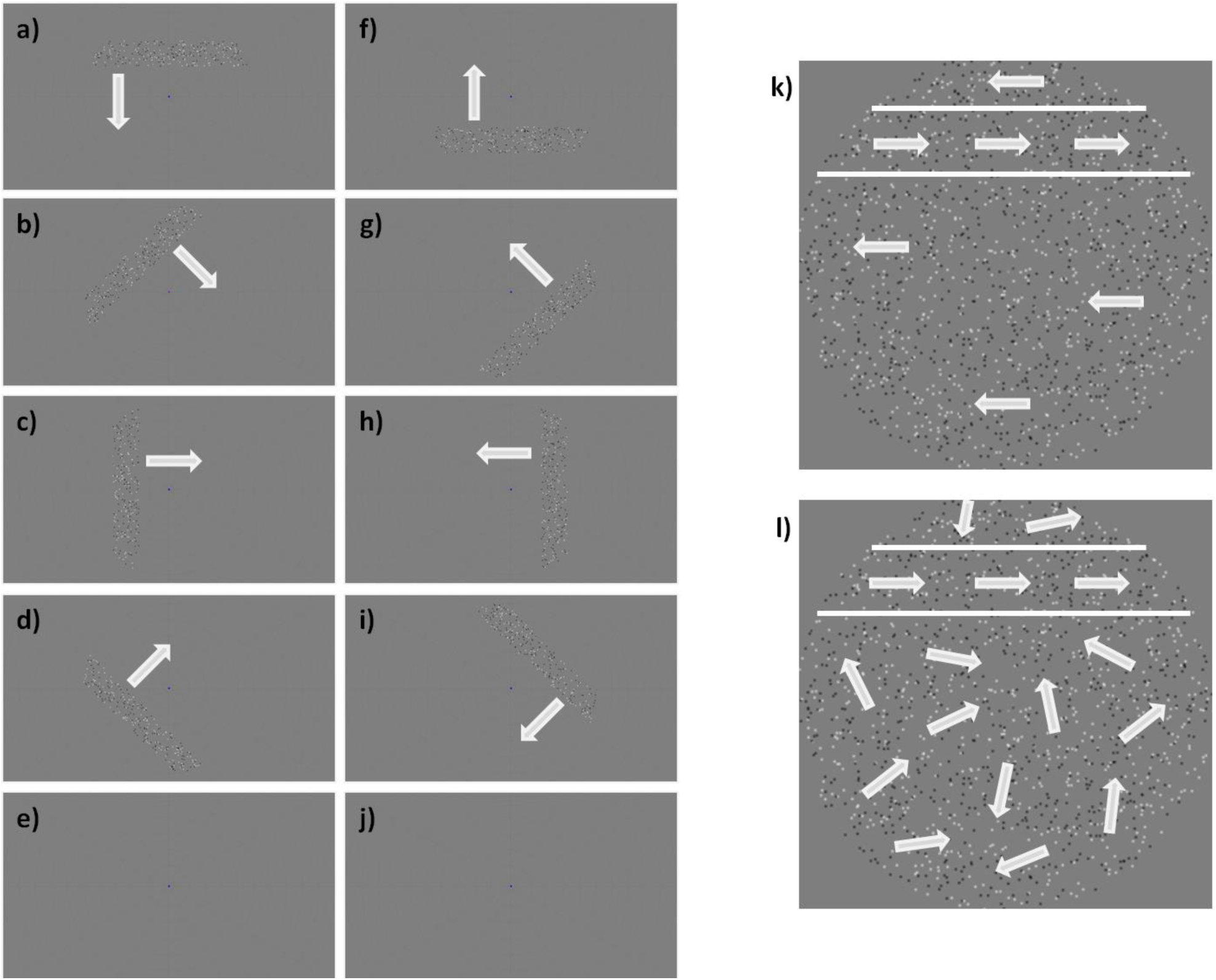
Schematic diagrams of the experimental set up in Experiment 1. (a) to (j) show one experimental run for the ‘bar-only’ condition, in the trial order presentation used for all stimuli in Experiment 1. Arrows indicate the direction of movement of the bar. Note in (b) that the fixation dot has changed colour, as part of the attentional task used in the experiment. (k) shows a schematic of the ‘kinetic’ condition. Within the bar (the area within the white lines) the dots moved in one direction (orthogonal to the bar movement); outside this area, they moved in the opposite direction. (l) shows a schematic of the ‘global’ condition; the movement inside the bar is in one direction, but outside the bar, the dots move in random directions.

Stimuli were created using MATLAB (MathWorks, Natick, MA) and the Psychophysics toolbox **(Brainard, 1997; Pelli, 1997)**. Stimuli were projected onto a screen (resolution 1920 × 1080 pixels, size 36.8 ×20.2cm) at the rear of the scanner bore, with the screen image reflected off a mirror attached above the head coil. The viewing distance was 67cm, meaning the screen subtended 30.7 × 17.1 degrees of visual angle on the retina. The refresh rate was 60Hz.

The experimental stimulus consisted of a field of 2000 dots (diameter = 0.09°) against a uniform median grey field. Half of the dots were randomly selected to be white, and the other half were black. The initial positions of the dots were randomly determined within a rectangular aperture 16 ×16° in size. A mask was then applied to the image, such that only the dots within a circle with 8° radius from the centre of the screen could be seen. A smaller circular mask (diameter = 0.8°) was also applied at the fixation point (diameter = 0.17°) to hide dots in this area and thus aid participants in maintaining their fixation. Further masks were also applied depending upon condition (see below). To further aid fixation, a low contrast “radar screen” pattern was shown behind stimuli (12 radial lines spaced 30 polar degrees apart, extending from just outside the fixation dot to the edge of the screen, (15.5°), along with 11 concentric rings centred on fixation increasing in radius in equal steps of 3°).

A ‘bar’ was defined as a strip within the circular stimulus region that was 2.4° wide and up to 16° in length (the total width of the hidden stimulus rectangle). This area was shifted over the course of each trial in 25 discrete steps of 0.7° and 1 second each, beginning at one edge of the stimulus aperture and ending on the opposite side. The bar could be rotated to start in any one of the four cardinal directions or the four oblique directions, giving eight different trial types. In one run, all eight different trial types were presented, along with two null trials where no bar was present. The null trials were always presented as the 5^th^ and 10^th^ trials. The order of the other directions was fixed for all participants and went anticlockwise from the first direction, which was where the bar was horizontal, starting at the top of the screen and moving downwards.

There were three different stimulus conditions that could be presented in a given run to the participants. In each condition, the dots within the bar always moved in the same way; what differed between conditions was the movement of the background dots outside the bar area.

In the ‘bar-only’ condition, only the dots within the bar were visible, and therefore the bar appeared to be moving across a grey background. In the ‘kinetic’ condition, the dots outside the bar (within the circular stimulus region) were always moving in the opposite direction to the dots within the bar, creating a ‘shearing’ effect which made the bar visible. In the ‘global’ condition, the dots outside the bar moved in random directions, allowing the bar to be detected as a 100% coherent global-motion stimulus against a background of noise in adjacent areas of the stimulus. While this condition therefore also contained kinetic boundaries, they were far less clear than the opposing directions used in the ‘kinetic’ stimulus. When a dot moved into the bar area (either through its normal progression or through a shift of the bar region), it started to move in the same coherent direction as all other bar dots. Similarly, when a dot left the bar area, it began to move randomly again. In all conditions, the null trials (with no bar present) had the same background motion as during the bar trials; a blank screen for the ‘bar-only’ condition, coherent motion in the ‘kinetic’ condition and random motion in the ‘global’ condition.

In all conditions, dots within the bar all moved in the same direction and moved along the length of the bar (so if the bar was moving from the top to the bottom of the screen, the dots moved from left to right or vice versa). All the dots (in both bar and background) changed direction by 180 degrees every 0.5 seconds, to prevent adaptation to one motion direction. Dots moved at 0.8°/second in all conditions. If any dots moved outside the aperture during the experiment, they were moved back one aperture width in the appropriate direction.

Each trial took 25 seconds, meaning that a run took 4 minutes and 10 seconds (plus a short period at the beginning of the run that was used to ensure that the fMRI signal had reached equilibrium). Each participant completed 4 runs for each condition, giving a total of 12 runs in the entire experiment. The order of the different conditions varied for different participants to control for order effects.

#### 2.1.3 Fixation task

Participants were instructed to focus on a blue fixation dot (diameter = 0.17°) at all times and to press a button on an MRI-compatible button box when they saw it change colour (to a red-purple). The probability of the blue dot changing colour was 0.01 every 200ms and the colour change periods lasted 200ms each. The results of this attentional task were unrecorded, and simply served to keep the participant fixated and alert throughout the experiment. An eye tracker (Eyelink 1000, sampling at the screen refresh rate of 60Hz) was used to monitor eye movements and ensure that participants were fixating correctly. We determined gaze stability using the methods outlined in **(Haas and Schwarzkopf, 2018)**; briefly, this involves calculating the median absolute deviation of the sampled gaze positions along both the horizontal and vertical dimensions for each run, and using these measures to compare the stability of gaze across conditions. Any run where fewer than 10 valid samples were taken was removed from further analysis. One participant was not eye tracked during either experiment, and therefore eye tracking data reflects the average of four participants in both Experiments 1 and 2. Analysis of the difference in eye position between conditions (both in the x and y directions) used general linear mixed models (using condition as a fixed factor, and subject and repeat number as random factors) followed by posthoc pairwise comparisons, with packages lme4 **(Bates et al., 2014)** and emmeans **(Russell, 2018)** in R (version 3.5.0).

#### 2.1.4 Data acquisition

Scans were acquired using a Siemens Avanto 1.5T MRI scanner with a 32-channel Siemens head coil located at the Birkbeck-UCL Centre for Neuroimaging. We used a modified version of the head coil without the eye visor to allow an unrestricted view of the screen, leaving 30 effective channels. We used functional T2*-weighted multiband 2D echo planar imaging with a multiband sequence (Breuer et al., 2005) and the following properties: voxel size = 2.3mm isotropic, field of view = 96 x 96, 36 slices, repetition time (TR) = 1s, echo time (TE) = 55ms, flip angle = 75°, and acceleration factor = 4. We collected 260-262 volumes (depending on stimulus condition) per run, and 4 runs were collected per condition for each participant. We also acquired a T1-weighted anatomical magnetisation-prepared rapid acquisition with gradient echo (MPRAGE) scan for each participant (TR = 2730ms, TE = 3.57ms) with a resolution of 1mm isotropic voxels.

#### 2.1.5 Analysis

The method used for analysing pRFs has been described previously **(Alvarez et al., 2015; Dumoulin and Wandell, 2008; Moutsiana et al., 2016; Schwarzkopf et al., 2014; van Dijk et al., 2016)**. In brief, the SamSrf MATLAB toolbox (available at http://dx.doi.org/10.6084/m9.figshare.1344765) models the pRF of each voxel as a 2D Gaussian in the visual field, incorporating a canonical haemodynamic response function based on the average of 26 participants in a previous study **(Haas et al., 2014)**. For each voxel the model finds the best-fitting visual field location, spread (standard deviation), and overall response amplitude of the pRF function.

Preprocessing of the fMRI data was carried out using SPM12 (Wellcome Centre for Human Neuroimaging, London, http://www.fil.ion.ucl.ac.uk/spm/software/spm12/). The first 10-12 volumes (depending on stimulus condition) were removed to allow the signal to reach equilibrium, leaving 250 volumes to be used in analysis for all participants and conditions. We then carried out intensity bias correction, realignment, unwarping and coregistration of the functional data to the structural scan, all using the default parameters built into the SPM software. FreeSurfer (https://surfer.nmr.mgh.harvard.edu/fswiki) was used to generate a 3D reconstruction of the grey-white matter surface **(Dale et al., 1999; Fischl et al., 1999)**, and the functional data was then projected to the cortical surface by finding the median position for each vertex of the surface reconstruction between the pial and grey-white matter boundary. Linear detrending was applied to the time series from each vertex in each run, and runs of the same stimulus condition were z-standardised and averaged together.

Population receptive field analysis was carried out on the occipital lobe data in a two-stage procedure. With a binary aperture describing the position of the bar element within each stimulus for each scanning volume (which was identical in each condition), we calculated its overlap with a profile of a pRF to predict the fMRI time series in the experiment. We first carried out a coarse grid search fit on data smoothed with a large kernel on the spherical surface (full width half maximum = 5), allowing calculation of the three pRF parameters that gave the maximal Pearson’s correlation between the predicted and observed time series for the full set of search grid parameters and vertices. These parameters were then used to seed an optimisation algorithm **(Lagarias et al., 1998; Nelder and Mead, 1965)** in a slow fine fit procedure on a vertex by vertex basis using unsmoothed data, allowing refinement of the parameter estimates and the calculation of an estimate of response strength.

Visual areas V1, V2 (dorsal and ventral), V3 (dorsal and ventral), V3A, V3B and MT+ (defined as TO1 and TO2; see supplementary materials for an example) were delineated based on reversals in the polar angle map from the ‘bar-only’ stimulus condition **(Sereno et al., 1995)**. For participants who only completed Experiment 2, the ‘transparent bar-only’ condition was used instead. These regions can be seen in Figure 2 and Figure 7. Throughout our main analyses, we used an R^2^ (goodness-of-fit) threshold of 0.05, which corresponds in our dataset to a p-value of 0.000367 (due to the number of observations per dataset and the number of free parameters in the pRF model). As our experimental conditions often show relatively weak and sparse responses, we chose this relatively liberal threshold to enable us to analyse the residual responses. However, we also carried out all analyses using a more conservative R^2^ value of 0.1, and results from these analyses can be seen in the supplementary material.

**Figure 2.**
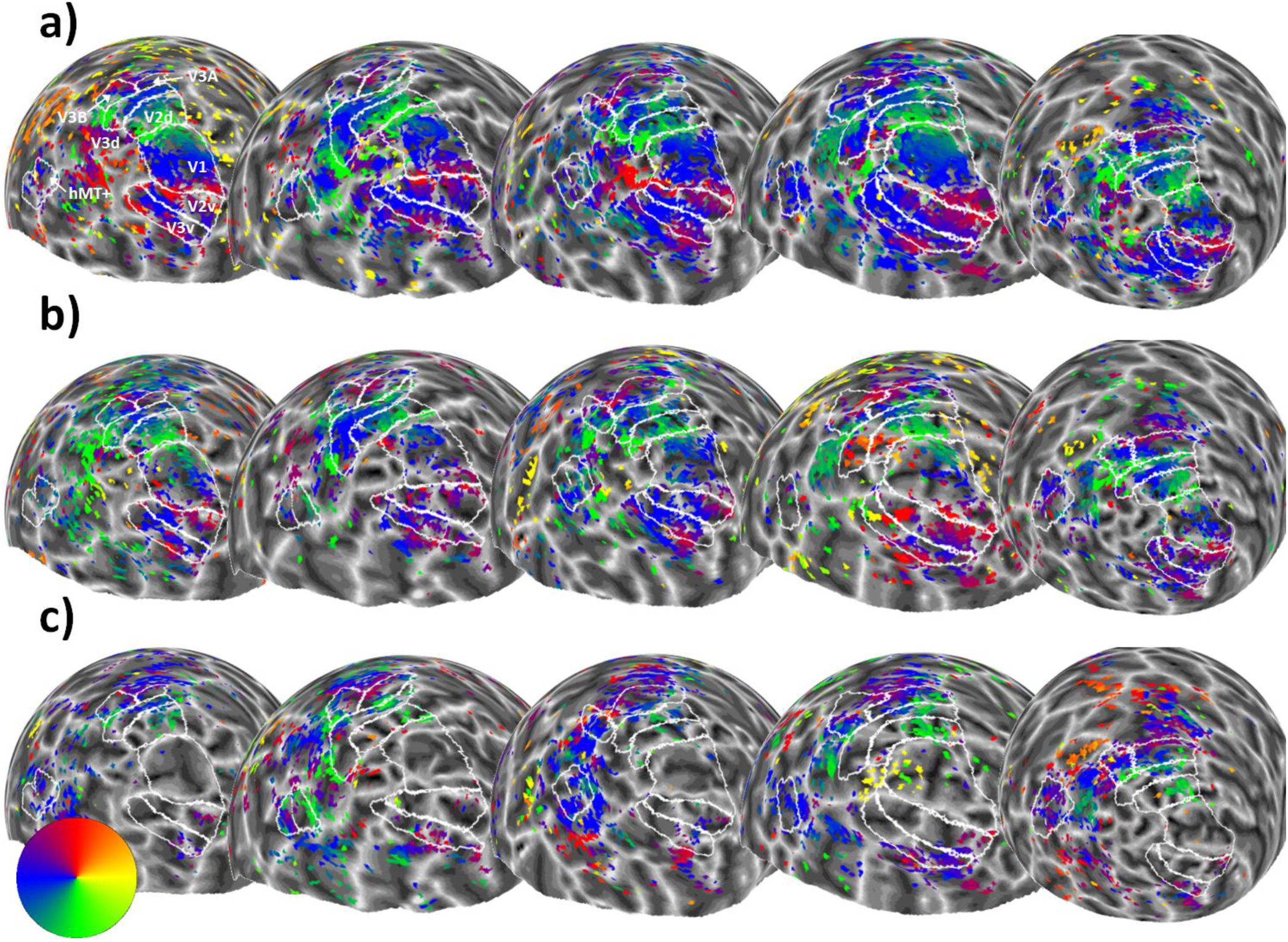
Sphere projection of polar angle data for the left hemispheres of all participants in Experiment 1. The colour of each vertex indicates the polar angle for the corresponding pRF centre (as indicated by the colour wheel). Each person’s data forms a column (subject 1 is on the far left, and subject 5 is on the far right), and stimulus condition forms a row. Manual delineations of visual areas V1, V2, V3, V3A, V3B and hMT+ (TO1/2) are shown. (**a**) Polar angle estimates for the ‘bar-only’ stimulus condition. (**b**) Polar angle estimates for the ‘kinetic’ stimulus condition. **(c)** Polar angle estimates for the ‘global’ stimulus condition.

### 2.2. Results and Discussion

#### 2.2.1 Relationships between maps

Figure 2 shows the left hemisphere polar angle maps for each experimental condition and each participant in Experiment 1 (R^2^ threshold = 0.05). Visual inspection of these images suggests that the ‘bar-only’ condition (Figure 2A) produces the clearest polar angle maps across participants, while the ‘kinetic’ condition (Figure 2B) tends to show a similar, if weaker pattern. The ‘global’ stimulus (Figure 2C) shows a weaker response still, with lower visual areas (e.g. V1) showing very little response. Although we had a relatively small number of participants, these general trends are highly consistent.

In order to quantify the consistency of these maps, we determined the correlation between the values for polar angle obtained for each vertex with each stimulus. As shown in Figure 3, the correlation between these polar angle estimates was overall clear, particularly for visual areas V2, V3, V3A and V3B where the average correlation was significantly different from zero. The correlation between conditions was less clear in V1 and MT+.

**Figure 3.**
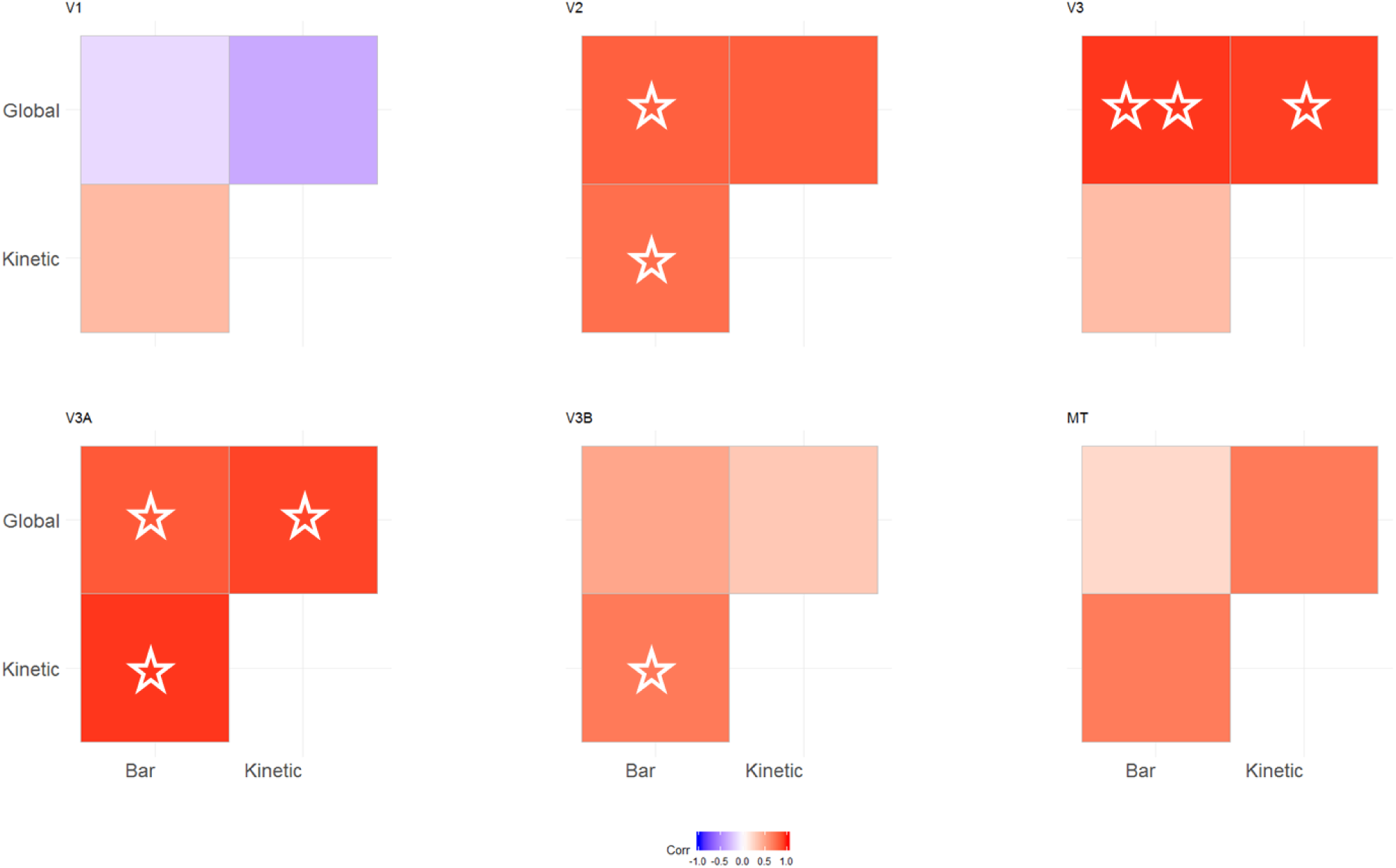
Correlation matrices comparing pRF polar angles between stimulus conditions in Experiment 1. The colour of each cell indicates the strength and sign of each vertex-wide correlation in polar angle. Circular correlations were calculated for each participant, then Z transformed and averaged across participants (as in (Haas and Schwarzkopf, 2018)). The symbols indicate whether the average correlation in individual cells is significantly different from zero (uncorrected). One star = p < 0.05. Two stars = p < 0.001.

We next determined the proportion of vertices within each of these visual areas responding retinotopically in the three experimental conditions (goodness of fit of the pRF model R^2^ > 0.05, Figure 4A). The bar stimulus produced the biggest response in areas V1-V3, which then dropped off for the higher visual areas (V3A, V3B and MT+). In contrast, responses to the kinetic stimulus increased across areas V1-V3, levelling off at V3A-V3B and then dropping in MT+. Responses to the global stimulus were even lower in the early visual areas, but again increased, reaching a peak at V3A and V3B. There were therefore large differences in stimulus responsivity in V1-V3, but these differences were much reduced in the higher visual areas. There was a significant interaction between condition and visual area in the final model of the data (interaction: χ^2^ = 336.06, p < 0.001; main effect of visual area: χ^2^ = 141.63, p < 0.001; main effect of condition: χ^2^ = 377.49, p < 0.001). This interaction also helps to explain the lack of significant correlation in polar angle values between conditions in area V1 (Figure 3) – although the bar stimulus produced clear polar angle estimates in a large number of V1 voxels, this was far less so for the other two stimuli.

**Figure 4.**
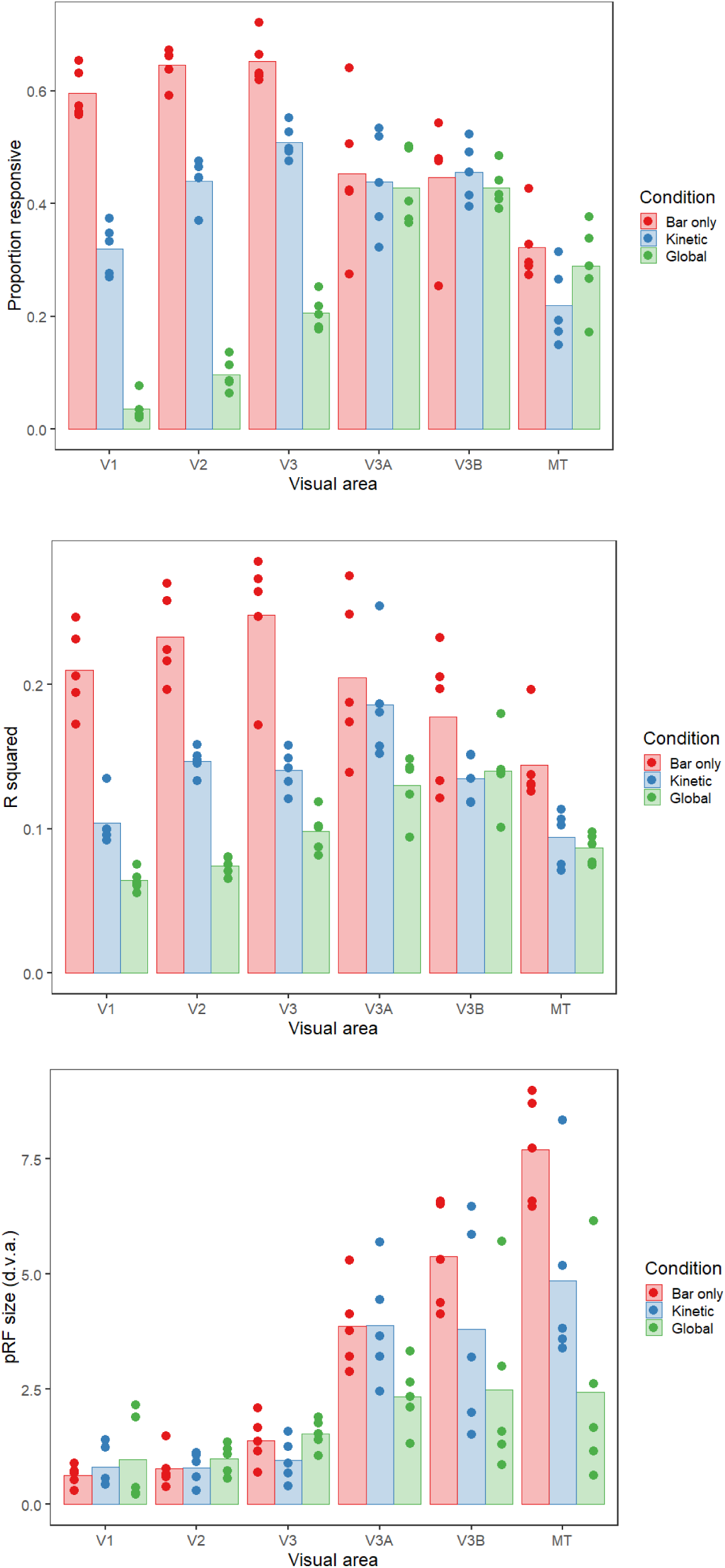
**(a)** Proportion of vertices responding, **(b)** goodness-of-fit and **(c)** pRF sizes for each condition and visual area in Experiment 1. The bars show the mean values across all subjects, and the points are individual data for each subject. In (a), this is the mean proportion of vertices responding for each subject, whereas for (b) and (c) these are the median goodness-of-fit values and pRF sizes respectively.

Comparing the goodness of fit across conditions (Figure 4B) showed a similar pattern, with initially large differences in R^2^ between conditions in the early visual areas that again decreased in higher regions. This interaction between condition and visual area was again significant (interaction: χ^2^ = 60.629, p < 0.001; main effect of visual area: χ^2^ = 62.747, p < 0.001; main effect of condition: χ^2^ = 242.912, p < 0.001). The area with the highest average R^2^ value also differed for each condition; the peak was in V3 for the bar stimulus, V3A for the kinetic stimulus and V3B for the global stimulus.

Finally, we analysed pRF size across conditions and visual areas (Figure 4C). Mean pRF sizes were smallest in the early visual areas, V1-V3, increasing in higher regions. pRFs were also relatively similar in early visual areas for all three conditions, but clear differences emerged in V3A, V3B and MT+. Here, pRFs were largest for the bar condition, smaller for the kinetic condition, and smaller again for the global condition (though in all cases larger than the equivalent condition in earlier regions). Again, there was a significant interaction between condition and visual area (interaction: χ^2^= 47.879, p < 0.001; main effect of visual area: χ^2^ = 174.505, p < 0.001; main effect of condition: χ^2^ = 24.434, p < 0.001). Similar results were seen when pRF size was examined as a function of eccentricity in the different brain areas and experimental conditions (see Supplementary Figure 2). pRF size was found to increase as a function of eccentricity in all brain areas, with the lowest rate of increase for the global condition.

#### 2.2.2 Control analyses

As there were clear differences in the proportion of responsive voxels in the three conditions, it is possible that the differences in pRF size between conditions were due to this reduction in the voxels included in each analysis, rather than a specific change in pRF size within each voxel. To examine this possibility, we analysed the data for the ‘bar-only’ condition using just the voxels that survived thresholding for the kinetic and global conditions (see Figure 5). The pattern of results is similar to Figure 4C – pRF sizes were again comparable in early visual areas for the three conditions, with clear reductions in pRF size for the kinetic and global conditions in areas V3A, V3B and MT+. In this case however, the reduction can be attributed to the differential selection of voxels responding to the same stimulus. In other words, the observed pRF size differences in Figure 4C are likely due to changes in the voxels that respond to these stimuli rather than active changes in pRF size across the different conditions. Using a linear model to compare the kinetic and global vertex conditions from the control analysis with the kinetic and global data in the original analysis showed no significant difference between the two data sets (χ^2^ = 0.758, p = 0.384).

**Figure 5.**
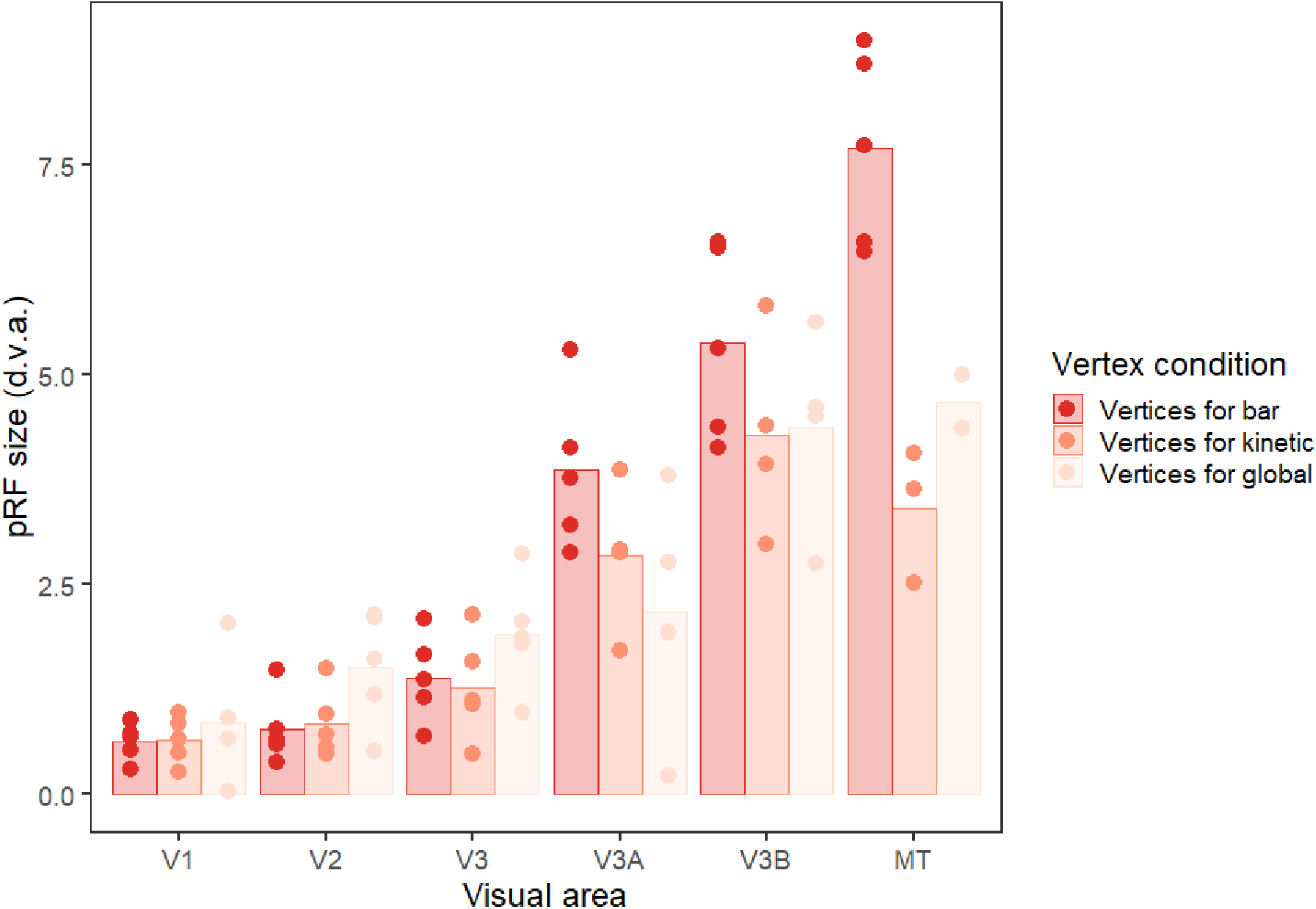
Plot to show pRF sizes for each visual area in Experiment 1 for the bar-only condition, using the responsive vertices for all three conditions. The bars show the mean values across all subjects, and the points are individual data for each subject (median pRF sizes). Any data points with a value of zero (obtained if the vertices for the condition did not overlap with any bar activation) were removed before plotting, leading to unequal numbers of data points in each condition.

It is also possible that differences between conditions could be attributed to differences in fixation stability between conditions. In a second control analysis we therefore examined the median absolute deviations of eye position, which on average were highly consistent and relatively low for both horizontal and vertical eye movements, averaging less than 0.5 degrees of visual angle for every condition (see Figure 6). General linear mixed models followed by posthoc pairwise comparisons suggested that there were no significant differences in eye position between conditions, either for the X or the Y direction (for X, bar-kinetic: t_30.14_ = 0.356, p = 0.933, bar-global: t_30.26_ = 0.457, p = 0.892, kinetic-global: t_30.35_ = 0.101, p = 0.994. For Y, bar-kinetic: t_30.35_ = −0.614, p = 0.814, bar-global: t_30.14_ = −0.503, p = 0.871, kinetic-global: t_30.43_ = 0.111, p = 0.993). Differences in fixation are therefore unlikely to have produced the above differences in pRF properties.

**Figure 6.**
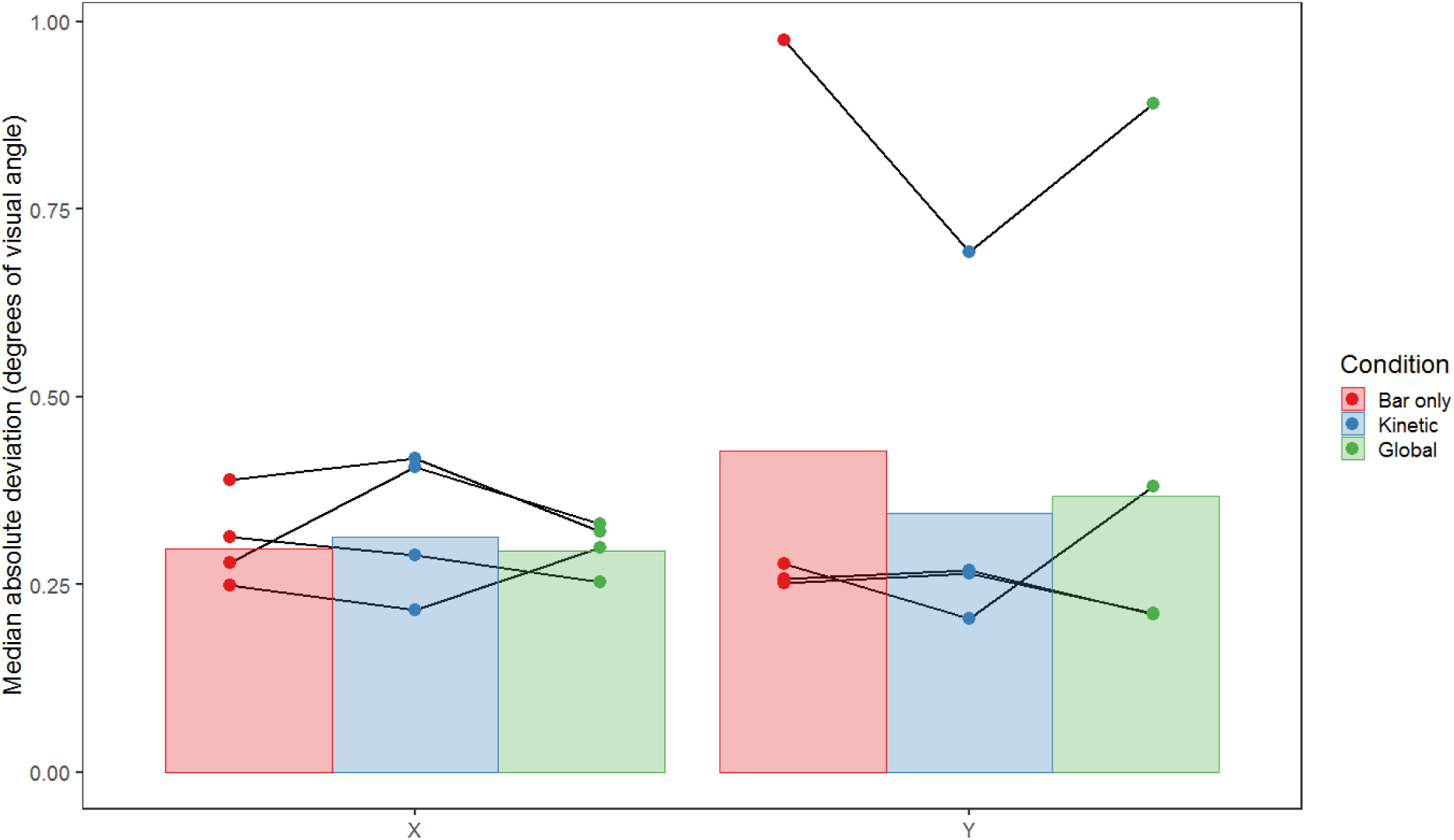
Plot showing the mean of the median absolute deviation of eye position across runs, conditions and observers (n = 4) for both the X (horizontal) and Y (vertical) dimensions of Experiment 1, in degrees of visual angle. Error bars are +/-1SD of the mean. Lines connect data points from individual subjects.

#### 2.2.3 Experiment 1 summary

Altogether, the three stimulus types (bar-only, kinetic, and global) produced clear differences in responsivity, goodness-of-fit, and pRF size across the visual hierarchy. Our control analyses reveal that these differences cannot be attributed to differences in gaze stability between the conditions and that the observed differences in pRF size are unlikely to reflect stimulus-driven changes in pRF size within each voxel. Rather, it seems likely that the observed differences in pRF size can be attributed to responses from different sub-populations of voxels in the three different conditions. We next turn our attention to the source of the differences in responsivity that appear to be driving these differences in pRF size. In particular, it is possible that differences in the visibility or salience of the bar between these conditions could drive the differences in responsivity. We explore this possibility in Experiment 2.

## 3. Experiment 2

As outlined in the introduction, estimates of pRF size and visual field location have been found to vary according to both the properties of the mapping stimulus and the attentional state of the observer. Although it is tempting to attribute the differences observed in Experiment 1 to differences in the effectiveness of these stimuli at driving the selectivity of various stages of the motion-processing hierarchy, the visibility of the bar element in each of our stimuli also varied between the three conditions. Most participants in Experiment 1 informally noted that the bar was less clear in the global condition than in the kinetic or bar-only conditions. It is possible then that the differences observed were driven by the visibility or salience of the bar, rather than any difference in the underlying selectivity of the motion detectors in each brain region.

In Experiment 2 we sought to test this by comparing the bar-only stimulus with two new stimuli. Firstly, we devised a second form of stimulus defined by global motion, a ‘transparent’ stimulus where the bar contained two sets of dots moving in opposite directions to give the appearance of two sheets moving transparently across each other **(Snowden and Verstraten, 1999)**. The global percept of these two directions requires an integration across space that is similar to that for a single direction of global motion amongst noise **(Edwards and Greenwood, 2005; Edwards and Nishida, 1999)**. Transparent motion is not perceived when dots with opposing directions are ‘locally balanced’ within small regions of the visual field **(Qian et al., 1994)**, perhaps due to an intermediate process of ‘local-motion pooling’ prior to the global-motion stage **(Edwards et al., 2012; Vidnyánszky et al., 2002)**. Although the responses of V1 neurons cannot distinguish between transparent and non-transparent stimuli, both fMRI **(Muckli et al., 2002)** and electrophysiological studies **(Qian and Andersen, 1994; Snowden et al., 1991)** show a differential response within MT/V5. Transparent-motion stimuli thus offer another stimulus with which we can assess whether pRF parameters differ when stimuli preferentially drive higher levels of the motion-processing hierarchy. We therefore constructed bar stimuli with two opposing directions of transparent motion within the bar, presented against a background of locally-balanced dots that do not appear transparent. If the differences in pRF parameters found in Experiment 1 are due to differences in motion selectivity, we would expect the transparent-motion bar stimulus to produce weaker responses in lower visual areas and a reduction in pRF size in higher visual areas, as with the ‘global’ condition.

Our second comparison stimulus was intended to examine the role of stimulus visibility in these effects. Transparent motion in particular has been found to be less visible in peripheral vision than in the fovea **(De Bruyn, 1997)**, which would likely create issues for the visibility of our transparent bar stimuli as they traverse the visual field, just as it may have been an issue in Experiment 1. We therefore compared these transparent stimuli with a stimulus bar that was not defined by differences in motion, but rather by a subtle difference in stimulus dot size. This bar stimulus would likely differentially drive the responses of early visual areas, given their potential role in the perception of object size **(Moutsiana et al., 2016; Murray et al., 2006; Pooresmaeili et al., 2013; Sperandio et al., 2012)**, but should not differentially drive the responses of higher motion-selective regions as effectively as the motion-defined bars used previously. The size difference in these stimuli does however lead to a substantial reduction in the visibility of the bar stimulus relative to the bar-only condition, particularly in peripheral vision. In particular, we selected a size difference that produced a similar level of subjective visibility to the ‘transparent’ condition (examined during pilot testing). Were this size-defined condition to produce similar responses to the motion-defined condition, this would suggest that visibility or salience is a more likely explanation for the observed differences than the stimulus selectivity of the underlying neural populations.

### 3.1. Materials and Methods

#### 3.1.1 Participants

Five participants (two male) took part in Experiment 2, including all four authors and one non-author participant from the first experiment (age range 28-39 years, mean age: 32.6 years). One participant was left handed. All had normal or corrected-to-normal visual acuity and provided written consent, as in Experiment 1.

#### 3.1.2 Stimuli

The second experiment was set up with the same apparatus and general stimulus properties as the first. Here there were three conditions related to the mapping stimuli: ‘bar-only’, ‘transparent’ and ‘size-defined’. The ‘bar-only’ condition in this experiment was identical to the ‘bar-only’ condition in Experiment 1, except that the dots were paired such that the two dots in each pair travelled in the same direction. However, different pairs travelled in opposite directions to give the impression of transparent motion with two ‘sheets’ of dots travelling over each other in opposite directions. As in the ‘bar-only’ condition of Experiment 1, the null trials for this condition contained only the uniform grey background. The speed of the dots was reduced in this condition to 0.34°/second to increase the impression of transparency.

In the ‘transparent’ condition, the bar was identical to the bar in the ‘bar-only’ condition. In this case however, the background also contained paired dots, with each dot in the pair moving in opposite directions, leading to the perception of non-transparent motion (as with ‘locally paired’ dot stimuli used previously; **(Qian et al., 1994)**. To keep dots within these local regions and avoid them unpairing over time, the motion of the dots periodically reversed. The timing of these reversals was randomised across dot pairs, so that reversals did not occur for all of the dots at the same time, a feature that enhances the percept of transparent sheets moving across one another **(Kanai et al., 2004)** when dot trajectories are limited (though not when locally paired, as in the background). The consistent pairing of dots between bar and background (differing only in their consistent vs opposing directions) meant that bar and background did not differ in dot density, and that the only feature to distinguish the bar was the percept of transparency against a background of non-transparent flicker.

The ‘size-defined’ condition contained a bar that was defined by a difference in dot size rather than by motion type; the dots in the bar were 0.10° in diameter against a background of dots with 0.09° diameter. All the dots in this condition moved in random directions, though with the same frequency of direction reversal as in the other two conditions (i.e. dots oscillated back-and-forth along a randomly selected axis). Null trials for this condition were the same as for the ‘global’ condition in Experiment 1.

To minimize anticipation effects in this experiment, the starting orientation and direction of the sequence (anticlockwise or clockwise shifts) was randomised for each condition and participant but kept constant across the four runs. Note that this meant that the movement direction of the bar always changed in a sequential fashion. All other presentation and analysis procedures were as in Experiment 1.

### 3.2. Results and Discussion

#### 3.2.1 Relationships between maps

As in Experiment 1, the bar-only stimulus produced clear and consistent polar maps across participants (Figure 7A). However, the transparent and size-defined stimuli produced much weaker and more variable maps (Figure 7B & C), particularly in lower visual areas (e.g. V1) where responses were considerably reduced. Interestingly, participants who self-reported that they were frequently unable to detect the transparency or size-defined bar stimuli (shown by red crosses in Figure 7) also had virtually no discernable map structure for these conditions.

**Figure 7.**
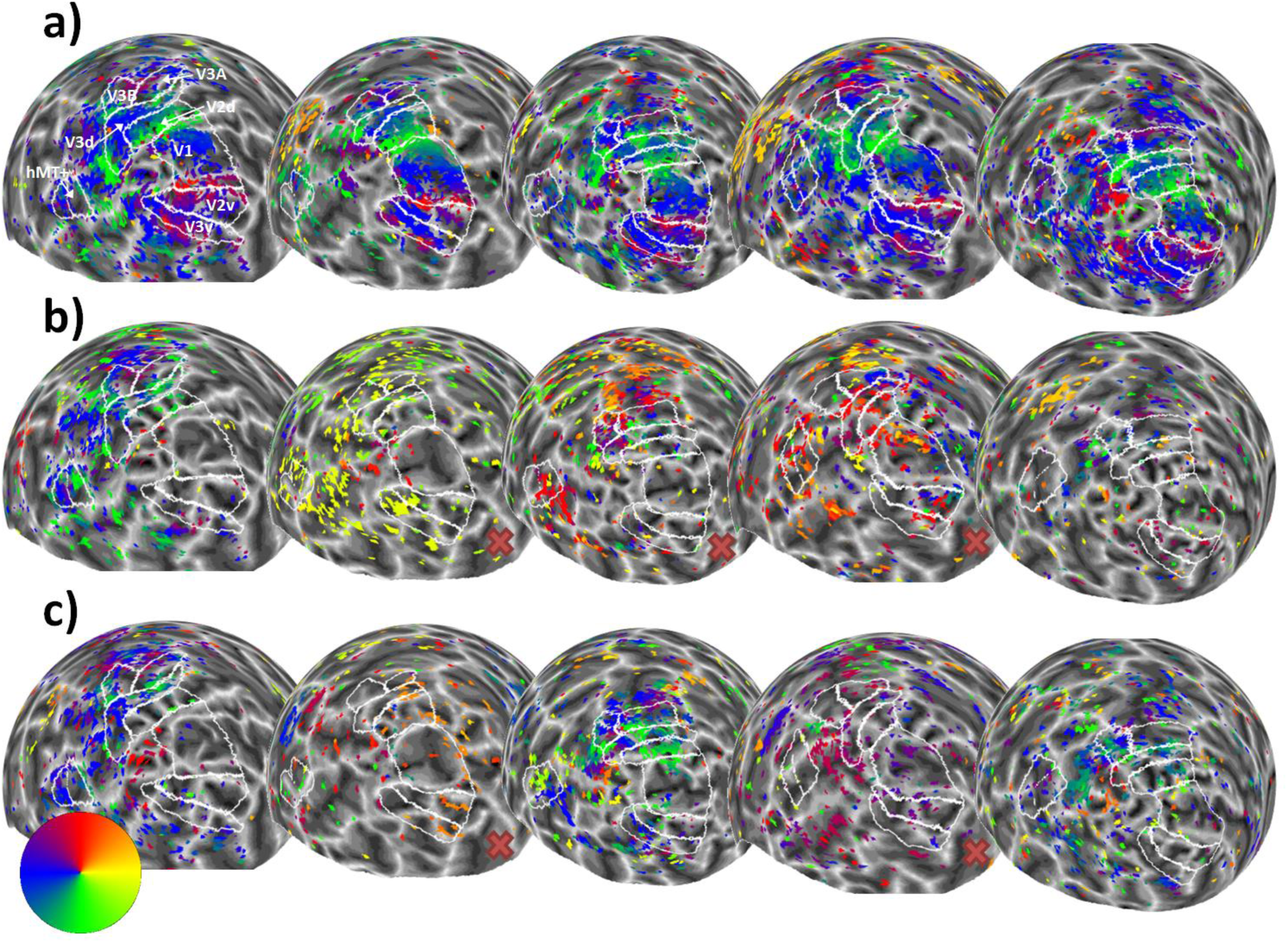
Sphere projection of polar angle data for the left hemispheres of all participants in Experiment 2. The colour of each vertex indicates the polar angle for the corresponding pRF centre (as indicated by the colour wheel). Each person’s data forms a column (subject 1 is on the far left and subject 5 is on the far right), and stimulus condition forms a row. Manual delineations of visual areas V1, V2, V3, V3A, V3B, and hMT+ (TO1/2) are shown (if the subject had taken part in Experiment 1, the delineations from this experiment were used). (**a**) Polar angle estimates for the ‘bar-only’ stimulus condition. (b) Polar angle estimates for the ‘transparent’ stimulus condition. (**c**) Polar angle estimates for the ‘size-defined’ stimulus condition. Red crosses indicate where participants self-reported low awareness of the stimulus when questioned after the experiment.

Correlations between these polar angle estimates in the 3 conditions were in general slightly weaker than in Experiment 1, but were still generally positive, with significant correlations overall only in areas V2 and V3A. As in Experiment 1, we next determined the proportion of vertices responding retinotopically in the different experimental conditions (see Figure 9A). Here, the bar-only condition produced similar response levels to the equivalent condition in Experiment 1, with these levels again decreasing in higher regions. In comparison, the transparent and size-defined conditions showed greatly reduced responsivity in early visual areas, similar to the global condition in Experiment 1. Responsivity in these conditions increased in later visual areas, though not quite to the level of the bar-only stimulus (unlike the global condition). There was a significant interaction between condition and visual area in the final model (interaction: χ^2^ = 52.330, p < 0.001; main effect of visual area: χ^2^ = 24.933, p < 0.001; main effect of condition: χ^2^ = 223.071, p < 0.001). This interaction is again likely to explain the lack of significant correlations in polar angle values between conditions, here given the clear drop in responsivity for the latter two stimulus types across the visual hierarchy.

**Figure 8.**
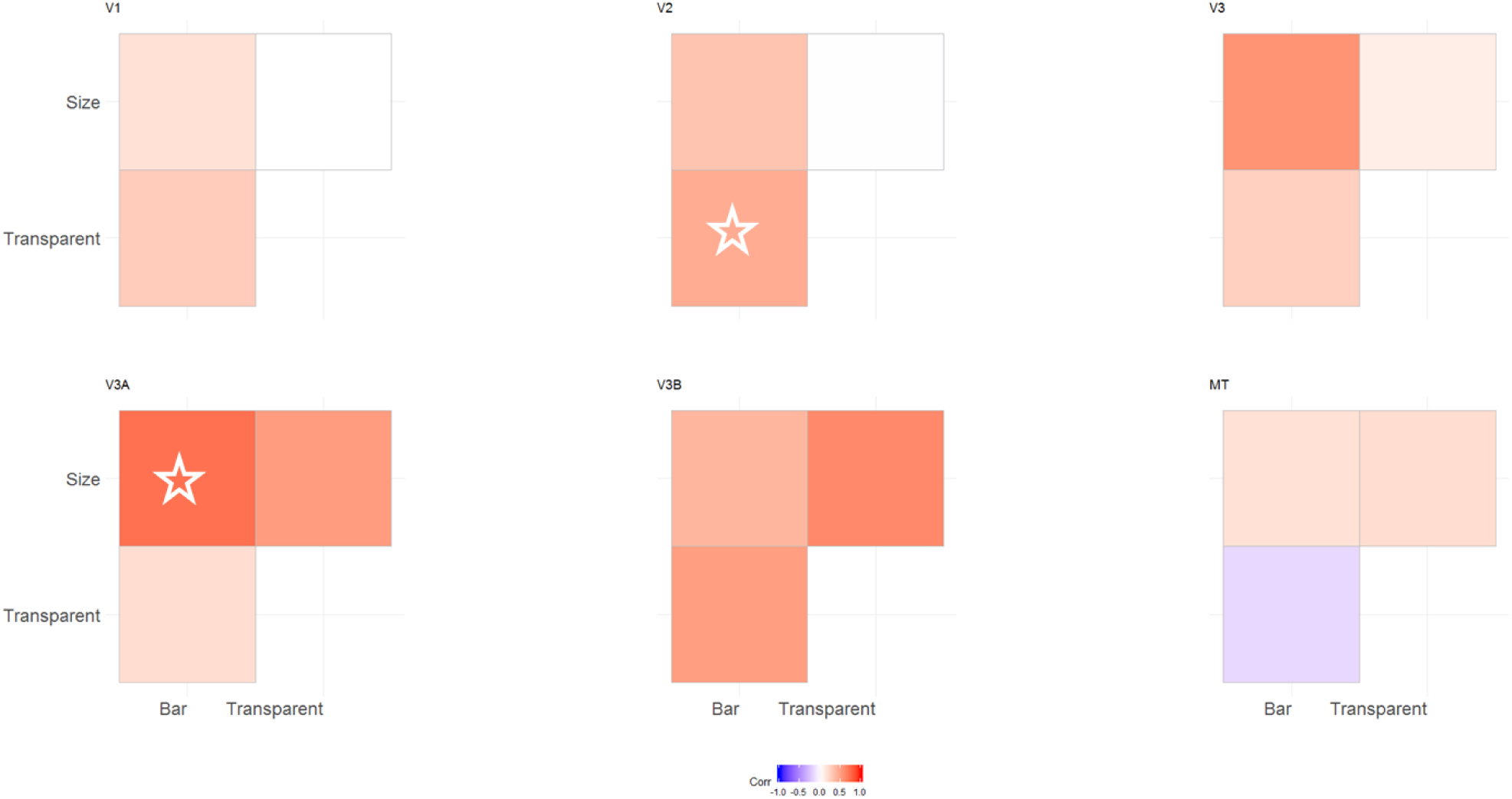
Correlation matrices comparing pRF polar angles between stimulus conditions in Experiment 2. The colour of each cell indicates the strength and sign of each vertex-wide correlation in polar angle. Circular correlations were calculated for each participant, then Z transformed and averaged across participants (as in (Haas and Schwarzkopf, 2018)). The symbols indicate whether the average correlation in individual cells is significantly different from zero (uncorrected). One star = p < 0.05. Two stars = p < 0.001. Note: one participant did not have enough data for valid correlations in the size-defined condition in V3B and MT, and so their data was not used for the average calculation).

**Figure 9.**
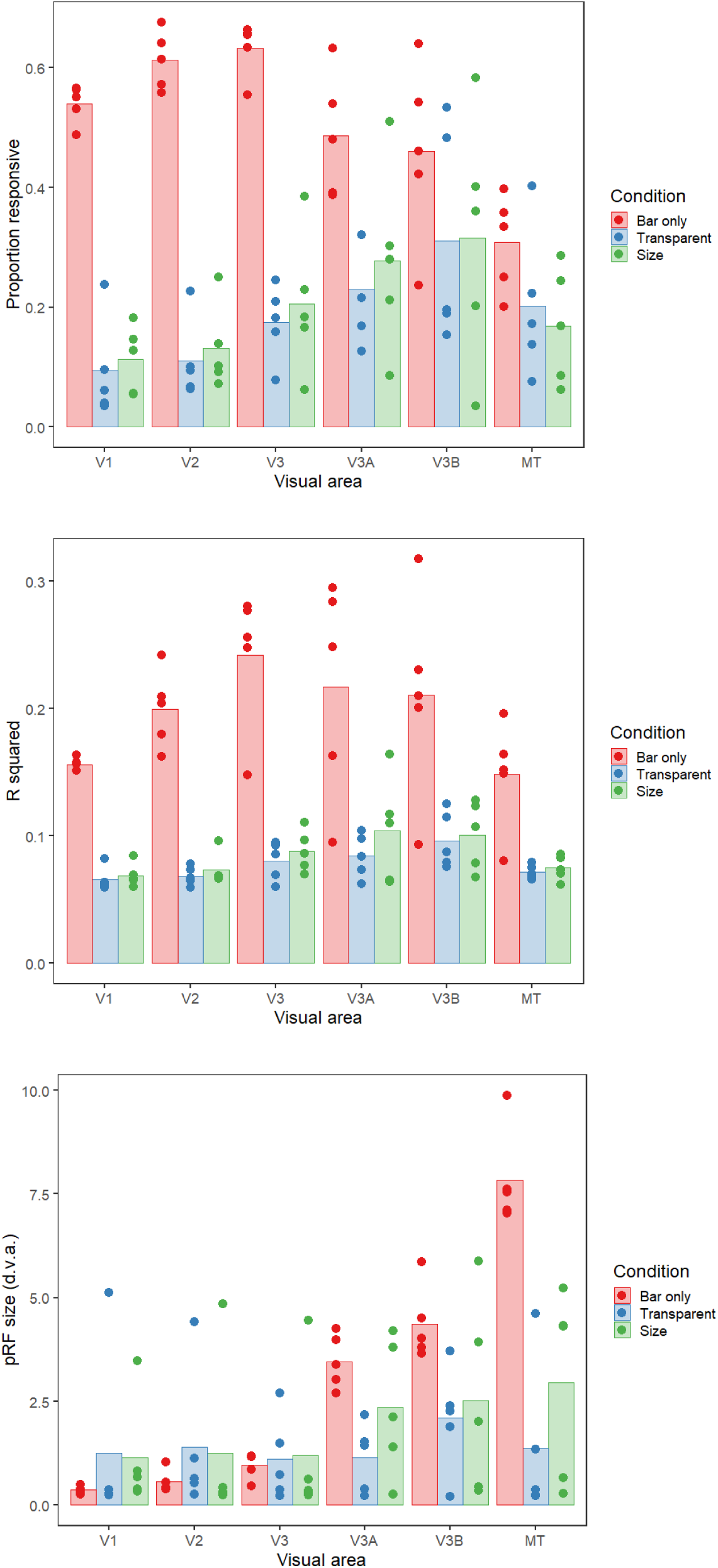
**(a)** Proportion of vertices responding, **(b)** goodness-of-fit and **(c)** pRF sizes for each condition (bar-only, transparent, and size-defined) and visual area in Experiment 2. The bars show the mean values across all subjects, and the points are individual data for each subject. In (a), this is the mean proportion of vertices responding for each subject, whereas for (b) and (c) these are the median goodness-of-fit values and pRF sizes respectively.

Goodness of fit also matched the pattern observed in Experiment 1, with the bar-only stimulus having much better goodness of fit compared to the transparent and size-defined stimuli. Goodness of fit was also worst in V1 and MT for all conditions, with R^2^ values increasing for more intermediate visual areas (Figure 9B). Modelling of the goodness of fit showed that there was no significant interaction between condition and visual area, (χ^2^ = 16.505, p = 0.086). However, there were significant main effects of visual area (χ^2^ = 30.060, p < 0.001) and condition (χ^2^ = 289.304, p < 0.001).

Finally, we considered differences in pRF size for the different conditions and visual areas (Figure 9C). Again, as in Experiment 1 there was a highly significant interaction between condition and visual area (interaction: χ^2^ = 64.806, p < 0.001; main effect of visual area: χ^2^ =72.031, p < 0.001; main effect of condition: χ^2^ = 21.572, p < 0.001). With the bar-only stimulus, pRF size increases dramatically across visual areas, with values comparable to those of Experiment 1. Although pRF values for the transparent and size-defined stimuli are comparable in early areas, the rate of increase is much lower than in the bar-only condition, resulting in considerably smaller pRFs in the highest areas. In MT these values were broadly comparable to the global condition of Experiment 1 for the size-defined stimulus, though generally much smaller for the transparent stimulus. pRF size also increased as a function of eccentricity in all different brain areas and experimental conditions, particularly for the bar-only stimulus (see Supplementary Figure 3).

#### 3.2.2 Control analyses

As in Experiment 1, a control analysis was run to examine whether the above differences in pRF size between conditions were due to this reduction in the voxels included in each analysis. We again analysed data for the bar-only condition using just the voxels that survived thresholding for the transparent and size-defined conditions. This again produced a clear reduction in pRF size for these two conditions in areas V3A, V3B and MT, suggesting that the observed differences in pRF size may be predominantly explained by differences in the responsivity of voxels (Figure 10). This is supported by statistical analysis suggesting that there is no significant difference between the transparent and size-defined conditions in the control analysis and in the original experimental analysis (χ^2^ = 0.111, p = 0.739).

**Figure 10.**
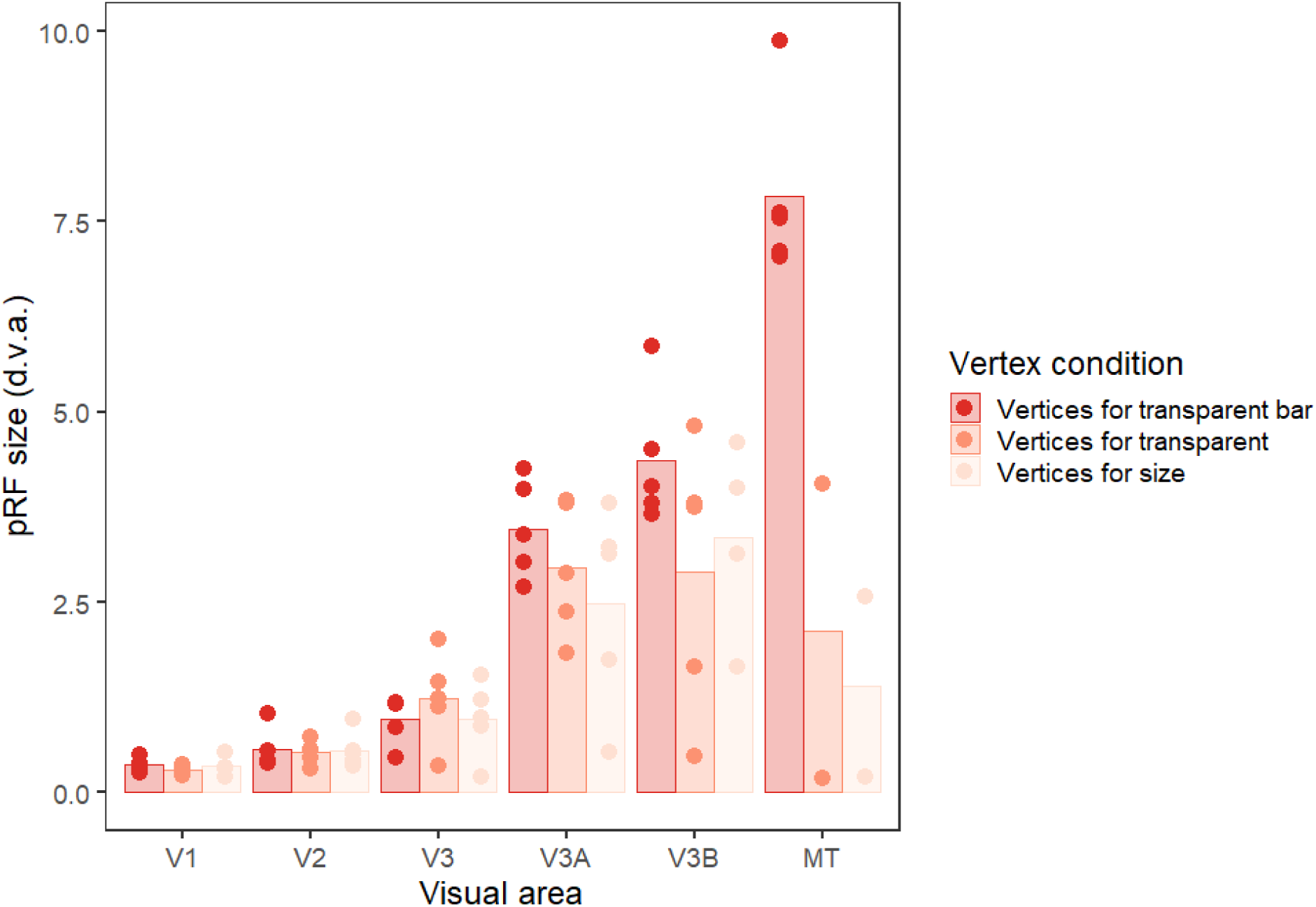
Plot to show pRF sizes for each condition and visual area in Experiment 2 for the transparent bar-only condition, using the responsive vertices for all three conditions. The bars show the mean values across all subjects, and the points are individual data for each subject (median pRF sizes). Any data points with a value of zero (obtained if the vertices for the condition do not overlap with any bar activation) were removed before plotting, leading to unequal numbers of data points in each condition.

As in Experiment 1, the results of Experiment 2 also do not seem to depend on eye movements, as the median absolute deviations of eye position were on average highly consistent and relatively low for both horizontal and vertical eye movements, with averages of less than 0.5 degrees of visual angle for both conditions (see Figure 11). General linear mixed models followed by posthoc pairwise comparisons suggested that there were no significant differences in eye position between the bar-only and size-defined conditions, either for the X or the Y direction (for X, bar-transparent: t_35.20_ = −0.055, p = 0.998, bar-size: t_35.04_ = −1..439, p = 0.332, transparent-size: t_35.20_ = −1..354, p = 0.375. For Y, bar-transparent: t_35.23_ = 0.667, p = 0.784, bar-size: t_35.05_ = −1..513, p = 0.297, transparent-size: t_35.23_ = −2.149, p = 0.094). As before, this suggests that participants were highly compliant with the fixation instructions and that differences in fixation stability cannot account for our results.

**Figure 11.**
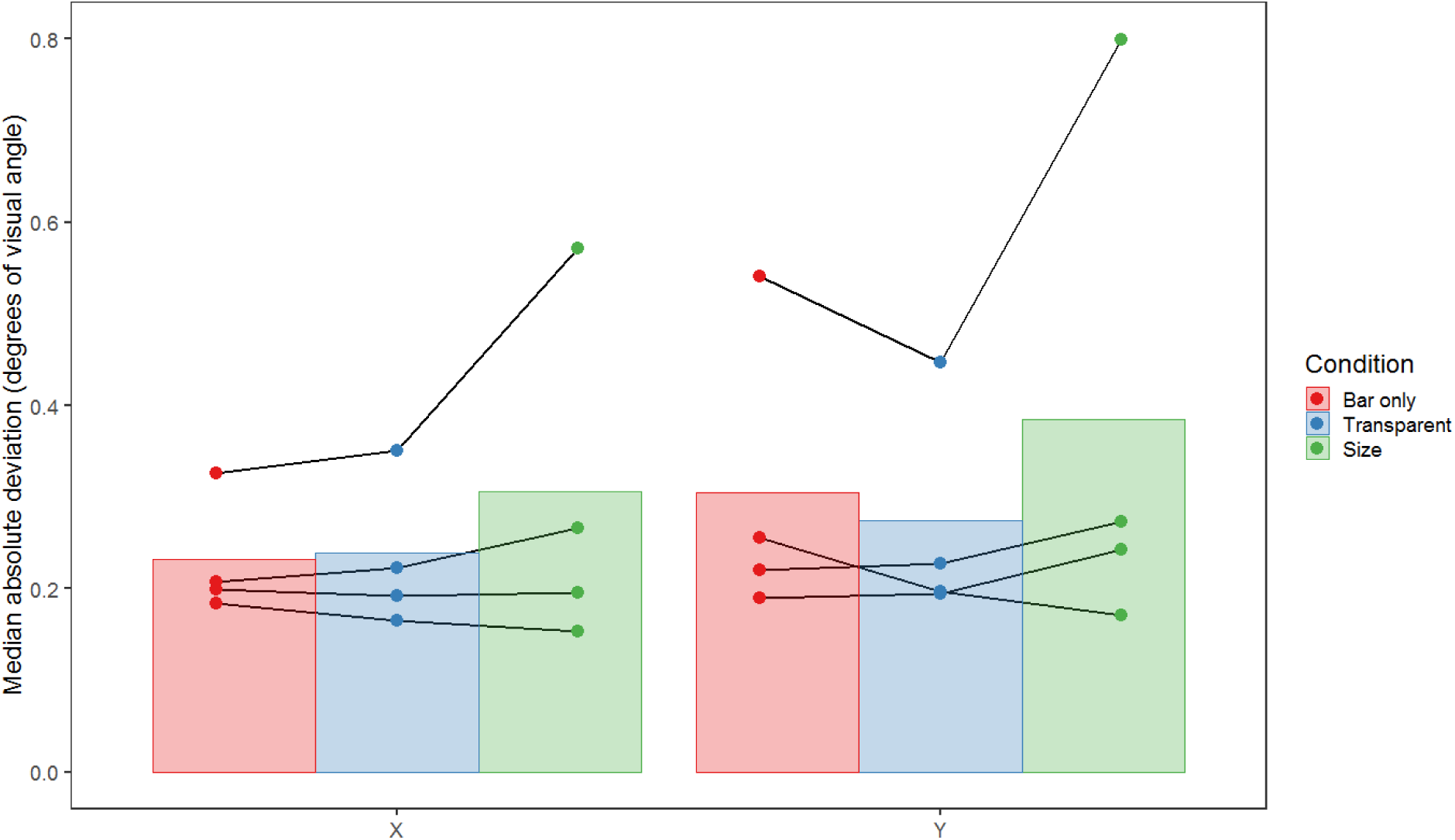
Plot showing the mean of the median absolute deviation of eye position across runs, conditions and observers (n = 4) for both the X (horizontal) and Y (vertical) dimensions of Experiment 2, in degrees of visual angle. Error bars are +/-1SD of the mean. Lines connect data points from individual subjects.

## 4. Experiment 3

In Experiment 2, we found very similar responses across visual areas for the transparent and size-defined stimuli, both of which differed from the bar-only stimulus. This indicates that the visibility or salience of the bar element could be important in determining responsivity and associated pRF properties. However, there are several possible aspects of visibility that could be involved. One is perceptual visibility, whereby the bar element may be less detectable (or more difficult to attend to) in some stimuli than others, particularly in peripheral vision. Another is neural ‘visibility’, where pRF analyses might be affected by the reduced signal-to-noise ratio in cases where there is a background signal (e.g. from the noise dots in the ‘global’ stimulus) as well as the responses to the bar element. To distinguish between these possibilities, we therefore carried out a control perceptual experiment, quantitatively assessing the visibility of the stimulus types used in Experiments 1 and 2 at different eccentricities, using psychophysical techniques. If perceptual visibility can explain the results found, we would predict that the ability of participants to detect the stimuli should follow the same pattern as the differences in responsiveness seen in Experiments 1 and 2. Specifically, bar stimuli should be highly visible, with a slight reduction in visibility for the kinetic stimuli, and further reductions for the global, transparent, and size-defined stimuli.

### 4.1. Materials and Methods

#### 4.1.1 Participants

Eight participants (three male) took part in the perceptual experiment, including two authors (who participated in both fMRI experiments), one non-author participant who took part in Experiment 1, and five naive participants (age range 21 – 37 years, mean age: 25.9 years). All had normal or corrected-to-normal visual acuity and provided written consent, as in previous experiments.

#### 4.1.2 Stimuli and procedures

The experiment was carried out in a laboratory setting (i.e. not in the scanner), with stimuli presented on a Display++ monitor (Cambridge Research Systems, UK) with a size of 71 × 39.5 cm, a resolution of 2560 × 1440 pixels, and a refresh rate of 120Hz. Viewing distance was 1m, with head movements restricted through the use of a chin and forehead rest. Responses were made via keypad. Stimulus parameters were set to subtend the same visual angle as in the fMRI set up.

On each trial, participants were presented with a single bar location of the stimuli used for the main fMRI experiments, meaning that the bar was presented in one of 22 different locations on the screen for 1 second (the middle three locations were excluded from this experiment to avoid ambiguity regarding their location relative to fixation). The participant’s task was to determine whether the bar was above, below, left or right of the fixation point, and press the corresponding button on a keypad to indicate their choice.

Each participant judged the bar location in five different types of stimuli: the three conditions used in Experiment 1 (bar only, kinetic and global) and the transparent and size-defined conditions from Experiment 2. All parameters used for these conditions were unchanged from the fMRI experiments. One block of the experiment used just one of the five stimulus types, with five repeats of all the possible positions (22) presented in two orientations (horizontal and vertical), giving 220 trials per block. The order of trials within a block was randomized. Each participant completed 10 blocks in total (two blocks of each stimulus type) and the order of these blocks was pseudorandomized (each stimulus type appeared once in the first five trials and once in the second five trials).

Testing was completed in two sessions of approximately 40-50 minutes each. At the beginning of the first session, naïve subjects were given practice trials for all stimuli. This involved showing the stimuli initially for 5s, and then running a practice block for each stimulus type where the bars were presented only at the innermost bar locations. This was done to ensure that naïve participants had a similar amount of experience with the stimuli as those who had participated in the fMRI experiments.

### 4.2. Results and Discussion

For each condition, responses were collated by eccentricity (collapsed across both visual hemifield and bar orientation) and scored as the proportion correct at each point. Figure 12 shows these responses along with the best-fitting linear function for each condition, a comparison that shows clear differences in visibility for the different stimulus conditions used in our experiments. Both the bar and kinetic stimuli were highly visible at even the furthest eccentricities. In contrast, the global stimulus was highly visible at central eccentricities, with a slight drop in visibility at 10-12 deg, although the difference in the slope of the linear fit was not significantly different from the bar condition (t = −1..473, p = 0.142). The size-defined stimulus was slightly harder to localise, even in central vision, with a steeper decline in visibility with eccentricity that was significantly different from that of the bar stimulus (t = −8.632, p < 0.001). The transparent-motion stimulus was even less visible at central eccentricities (though performance was still well above chance), with a similarly steep decline in visibility with increasing eccentricity that was again significantly different from the bar stimulus (t = −7.857, p < 0.001). Detectability of the size-defined and transparent-motion stimuli was more variable across participants (as indicated by the larger error bars), similar to the variation in visibility reported in Experiment 2.

**Figure 12.**
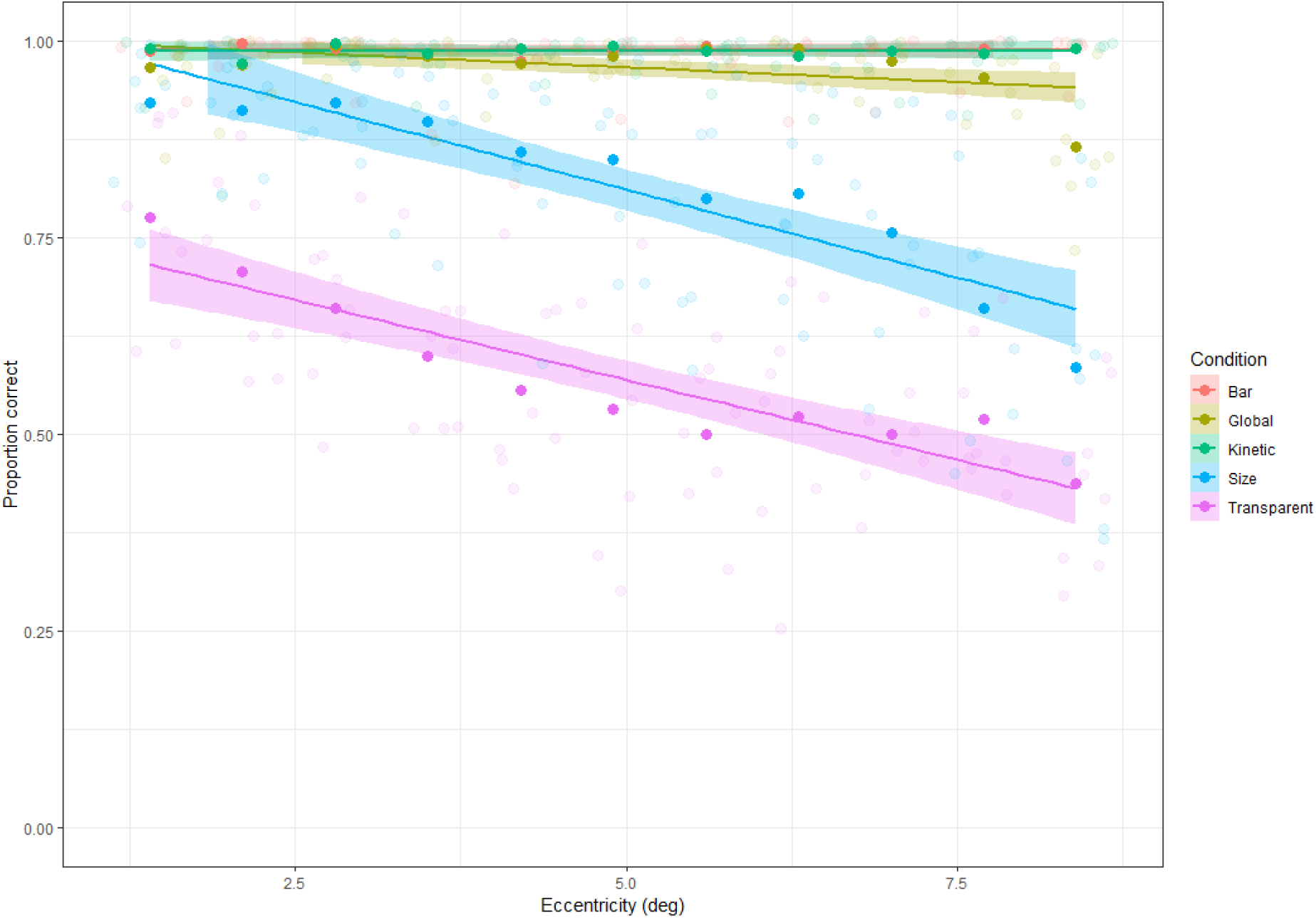
Plot showing the mean percent correct (n=8 participants) for each of the five stimulus types in the perceptual experiment, across different eccentricities. All individual data points are represented, along with a linear fit to the full data set for each condition.

Overall however, the above pattern of visibility does not closely match the variations observed in the properties of the pRFs measured with these stimuli. In particular, the global condition was similarly visible to the kinetic and bar conditions but showed a very different pattern of pRF data in Experiment 1. Conversely, the size-defined and transparent conditions showed clear reductions in visibility relative to both the global condition and to each other, and yet the pRF properties measured with these stimuli were both similar to those of the global condition. We conclude that stimulus visibility is unlikely to account for these differences in pRF properties.

## 5. Discussion

In this study, we show that retinotopic mapping stimuli defined by motion produce clear and highly consistent differences in the properties of population receptive fields (pRFs) measured across the visual hierarchy, including responsivity, goodness of fit, and pRF size. As predicted, we show that a bar mapping stimulus defined by moving dots (against a blank background) produces strong pRF maps in early visual areas, with responsivity decreasing and pRF sizes increasing in areas higher in the motion processing hierarchy. More complex motion stimuli, such as bars defined by kinetic and global motion (against backgrounds of opposing motion or noise, respectively), produce a much lower degree of responsivity in early visual areas, with a reduction in pRF sizes for higher visual areas, and reductions in goodness of fit across the hierarchy as a whole. Control analyses further suggest that the reduction in pRF size can be attributed to the reduction in the voxels included for each stimulus, rather than changes in pRF size within voxels. Although it is tempting to attribute this to differences in the potential for these visual areas to distinguish these higher-order stimuli, a second experiment showed highly similar patterns of responsivity, goodness of fit, and pRF size across visual areas for a stimulus defined by transparent motion (against a non-transparent background) and a size-defined stimulus (with no differences in motion) that were reduced in visibility. This suggests that the observed differences in pRF properties are not specific to mapping stimuli defined by differences in stimulus motion.

As outlined in the introduction, evidence from a variety of experimental approaches suggests that motion is processed hierarchically in the visual system, with local motion processed in early visual areas, such as V1, and global motion processed in higher areas, such as MT+ **(Adelson and Movshon, 1982; Braddick et al., 2001; Movshon, 1986; Van Essen and Gallant, 1994; Williams and Sekuler, 1984)**. These differences in selectivity predict differences in pRF size and responsivity between local motion defined stimuli (such as our bar stimulus) and global motion defined stimuli (such as our global stimulus) for these different areas. Our results indeed show these differences, sometimes quite strikingly; the lack of response in V1 for the global stimulus is highly consistent across observers. Support for differences derived from other forms of motion selectivity is less clear. In particular, it has been suggested that kinetic boundary stimuli are processed preferentially in visual area V3B **(Van Oostende et al., 1997)**. However, our results show that responses were equally as strong in V3A, with comparable pRF sizes and goodness-of-fit values that were, if anything, better than V3B. Our results therefore suggest that although there were clear differences in the pRF properties measured in early visual areas with these stimuli, amongst higher areas these kinetic boundary stimuli produced widespread changes in pRF properties that are difficult to localize to any one area, supporting the notion that motion boundaries may be processed in a wide number of areas in the visual cortex **(Larsson et al., 2010; Larsson and Heeger, 2006)**.

The fact that we see very similar response patterns for both our ‘transparent’ and ‘size-defined’ conditions in Experiment 2 further suggests that we should be cautious about attributing our results to differential motion processing in distinct visual areas and should instead consider alternative explanations. One possibility is that the results could reflect a decrease in visibility because of the presentation of these bar mapping stimuli in peripheral vision. Our sensitivity to global motion **(Raymond, 1994)**, transparent motion **(De Bruyn, 1997)**, and kinetic boundaries or motion-defined form **(Regan and Beverley, 1984)** are all known to decline in the periphery. These declines can be corrected for by adjusting the contrast, size, and speed of stimuli **(Hess and Aaen-Stockdale, 2008; Regan and Hamstra, 1991)**, at least to some extent. However, were the lack of eccentricity scaling a problem with our stimuli, we should have seen an increase in average pRF size with our motion-defined stimuli due to the loss of responsivity from neurons with small receptive fields (as seen in comparisons between size-invariant and eccentricity-scaled bar stimuli; **(Alvarez et al., 2015)**. Our psychophysical results in Experiment 3 also show that while there are decreases in visibility in peripheral vision, these do not show a consistent relationship with our pRF data. For instance, while the global, transparent and size-defined conditions all showed reductions in pRF responsivity relative to the bar-only stimulus, the bar element in the global condition was in fact clearly visible at all but the most extreme eccentricities. Conversely, the size-defined bars were much harder to localise, with further reductions for the transparent stimulus, and yet pRF responsivity rates for these two conditions were similar. This suggests that perceptual visibility is not the main factor driving the responses we observe.

Previous research has shown that differences in attention to peripheral stimuli can also influence neural responses. One study found decreased responses in V1 (but stronger responses in parietal and frontal areas) when participants distinguished between stimulus and background entirely attentionally **(Saygin and Sereno, 2008)**. It has also recently been shown that it is possible to map retinotopic responses for bar stimuli defined by illusory contours, occluded parts of a bar, or very low luminance contrast **(Haas and Schwarzkopf, 2018)**, suggesting that the mapping reflected spatial attention rather than specific visual properties of these stimuli. In the current experiment, it is likely that the more complex second-order stimuli used were more attentionally demanding than the ‘bar-only’ stimuli. Along these lines, second-order motion has been found to be more difficult to process at multiple locations compared to first-order motion, suggesting that second-order motion is more attentionally demanding **(Lu et al., 2000)**. Direction discrimination thresholds for second-order motion are also influenced more strongly by attention than thresholds for first-order motion **(Allen and Ledgeway, 2003)**. It may be then that the greater attentional demands required to detect second-order stimuli (like our bars defined by differences in global motion) are more important for determining the responsivity and properties than the stimulus parameters. Again, however, the results of our psychophysical experiment argue against this – the global stimulus in particular was highly visible across the visual field (suggesting that observers had no difficulty attending to these bar elements) and yet pRF responses were similar to those for the transparent stimulus that was much less visible (and which therefore may have presented difficulties for attention). In other words, attention does not seem to offer a complete explanation for our results.

Another possibility is that our motion-and size-defined stimuli may have produced an illusory sense of depth for the bar stimulus, which may then have altered our measured pRF properties. It is known that areas such as V3B and V3A are involved in the processing of depth cues **(Tyler et al., 2006)**, and particularly with integration of depth cues with other signals, such as motion **(Ban et al., 2012)**. However, we think this is unlikely to be a complete explanation of our results, as participants did not report strong depth percepts for any of our stimuli. In addition, it is not clear that this hypothesis would explain the patterns observed in our results; we did not see markedly stronger responses in V3A/V3B, as has been observed in previous fMRI studies of depth perception **(Anzai and DeAngelis, 2010; Backus et al., 2001; Tsao et al., 2003)**. A related suggestion is that the responses in higher visual areas may be a consequence of surface segmentation cues; however, again, research has shown that early visual areas such as V1 are also activated by texture detection and surface segregation processes **(Scholte et al., 2008)**.

Finally, rather than psychophysical visibility, it is likely that the visibility of the neural responses to mapping stimuli (relative to background activity and noise levels) may be an issue in the measurement of population receptive fields. pRF analyses rely on a difference in BOLD response when the stimulus bar and the background are presented in a given location of the visual field (though of course the BOLD response is an indirect measure of differences in neural processing that may reflect vascular responses in some situations; **(Logothetis, 2008; O’Herron et al., 2016)**. Changes in the response to the background stimulus may therefore affect our ability to derive these measures. In particular, although unidirectional global motion typically drives BOLD responses in MT to a greater extent than incoherent noise **(Braddick et al., 2001)**, incoherent noise stimuli still produce an increased response within MT relative to stationary stimuli **(McKeefry et al., 1997)**. The same would be true for our kinetic boundary and transparent stimuli. It is possible therefore that the luminance-defined differences produced by the bar-only stimulus produce a clearer difference between bar and background responses than the motion-defined differences of the other conditions, consistent with the observed reductions in goodness of fit for the pRF parameters derived using these motion-based stimuli. This could also explain why size-defined stimuli produced a similar pattern of results, given that the noise dots used in these stimuli would similarly decrease the difference in BOLD response to the stimulus bar relative to the background. Larger pRF estimates are likely to be particularly vulnerable to this issue, given that these voxels tend to show the worst goodness-of-fit. For instance, in Experiment 1 there was a clear negative correlation between R^2^ and pRF size in V1, even with the bar-only stimuli (ρ = −0.721, p < 0.001). Voxels with large pRF estimates may thus be the first to drop out with our motion-based stimuli, leading to our observed reductions in pRF size. Were this to be the case, our findings would in fact reflect the selectivity of visual brain regions for motion (given the increased responsivity to the stimulus background), though the reduction in pRFs could not be strictly interpreted as a property of the underlying neural populations. Given that our behavioural data suggests that simple psychophysical visibility is not well matched to our pRF results, we would argue that this explanation provides the most parsimonious explanation of our results.

Previous work has suggested that when the stimulus bar is defined by orientation contrast, reductions in pRF size in higher visual areas (such as LO1 and LO2) may be caused by the stimulus mainly driving neurons sensitive to orientation contrast **(Yildirim et al., 2018)**. Here, while we also find reductions in pRF size for our more complex second-order conditions, the similarity in responses between very different conditions (like the size-defined and transparent stimuli) leads us to argue that this reduction can be more parsimoniously explained by reductions in the signal-to-noise ratio of the neural signal, as discussed above. Of course, this does not mean that second-order stimuli are not useful (for example, they may potentially improve the accuracy of pRF estimates by reducing BOLD displacement; **(Olman et al., 2007; Yildirim et al., 2018)**, and it does not mean that there may not be stimulus-specific signals in pRF mapping; for example, recent studies have shown that varying the orientation or direction of motion of the carrier stimulus within the bar apertures used for mapping can lead to differences in pRF parameters **(Dumoulin et al., 2014; Harvey and Dumoulin, 2016)**. We simply urge future researchers to be cautious when interpreting the functional meaning of changes in pRF properties.

In conclusion, we find evidence for variations in the properties of retinotopic maps for different motion-based stimuli. In particular, we find clear retinotopic maps for stimuli defined by a moving bar of dots against a blank background, but much weaker maps when the bar was defined by coherently moving dots against a background of either incoherent or oppositely-moving dots, or by transparent compared to non-transparent motion. However, the similar maps derived from stimuli defined by size differences suggest that these differences do not reflect a change in the responsivity of neurons in different visual areas to different motion properties. We similarly rule out variations in perceptual visibility or attentional selection of the bars with our behavioural data. Rather, we suggest that it is the visibility of the neural signal for retinotopic mapping stimuli, as defined by the signal-to-noise ratio between bar and background responses, that is the most important driver of pRF properties.

## Supplementary Materials

Video S1: Example bar-only stimulus, Video S2: Example kinetic stimulus, Video S3: Example global stimulus, Video S4: Example transparent bar-only stimulus, Video S5: Example transparent motion stimulus, Video S6: Example size-defined stimulus. Supplementary document: containing an example delineation of TO1 and TO2, eccentricity plots and analyses using a R^2^ threshold of 0.1.

## Acknowledgments

Supported by ERC Starting Grant WMOSPOTWU to DSS, and Career Development Award MR/K024817/1 from the UK Medical Research Council to JAG. Our thanks to Jingxiu Cheng for his help with data collection in the behavioural experiment.

## Author Contributions

All authors conceived and designed the experiments and performed the experiments; A.H. and D.S.S. analysed the data; D.S.S. contributed analysis tools; A.H. wrote the paper; all authors edited the paper and approved the final version.

## Conflicts of Interest

The authors declare no conflict of interest. The funding sponsors had no role in the design of the study; in the collection, analyses, or interpretation of data; in the writing of the manuscript, and in the decision to publish the results.

## Supplementary material

### Delineation of TO1 and TO2

**Figure S1:**
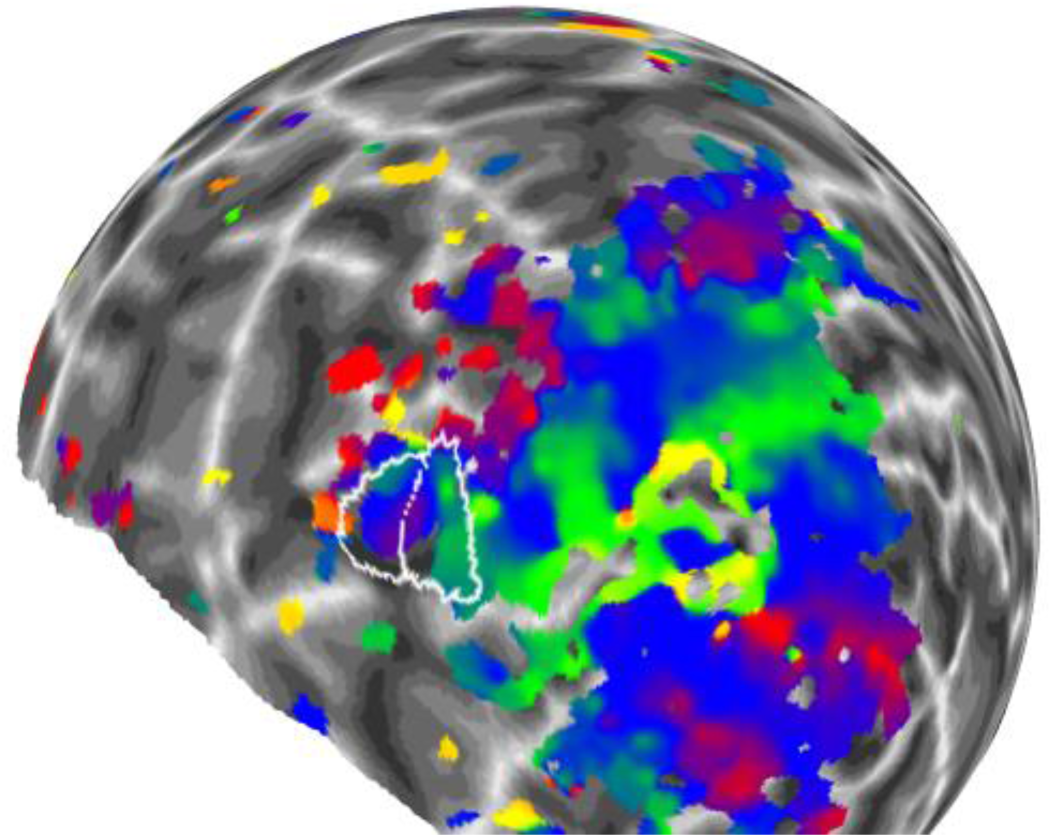
An example delineation of areas TO1 and TO2 on the smoothed retinotopic maps for a single participant (using data from the dot-only bars in Experiment 1, and a threshold of 0.06). The border between them was determined as being an upper vertical meridian reversal (red), and the outer borders on either side were either a horizontal (blue) to lower vertical meridian (green). For analyses however, we combine TO1 and TO2 into one MT+, so the exact distinction is not relevant for analysis.

#### Plots of pRF size against eccentricity

**Figure S2:**
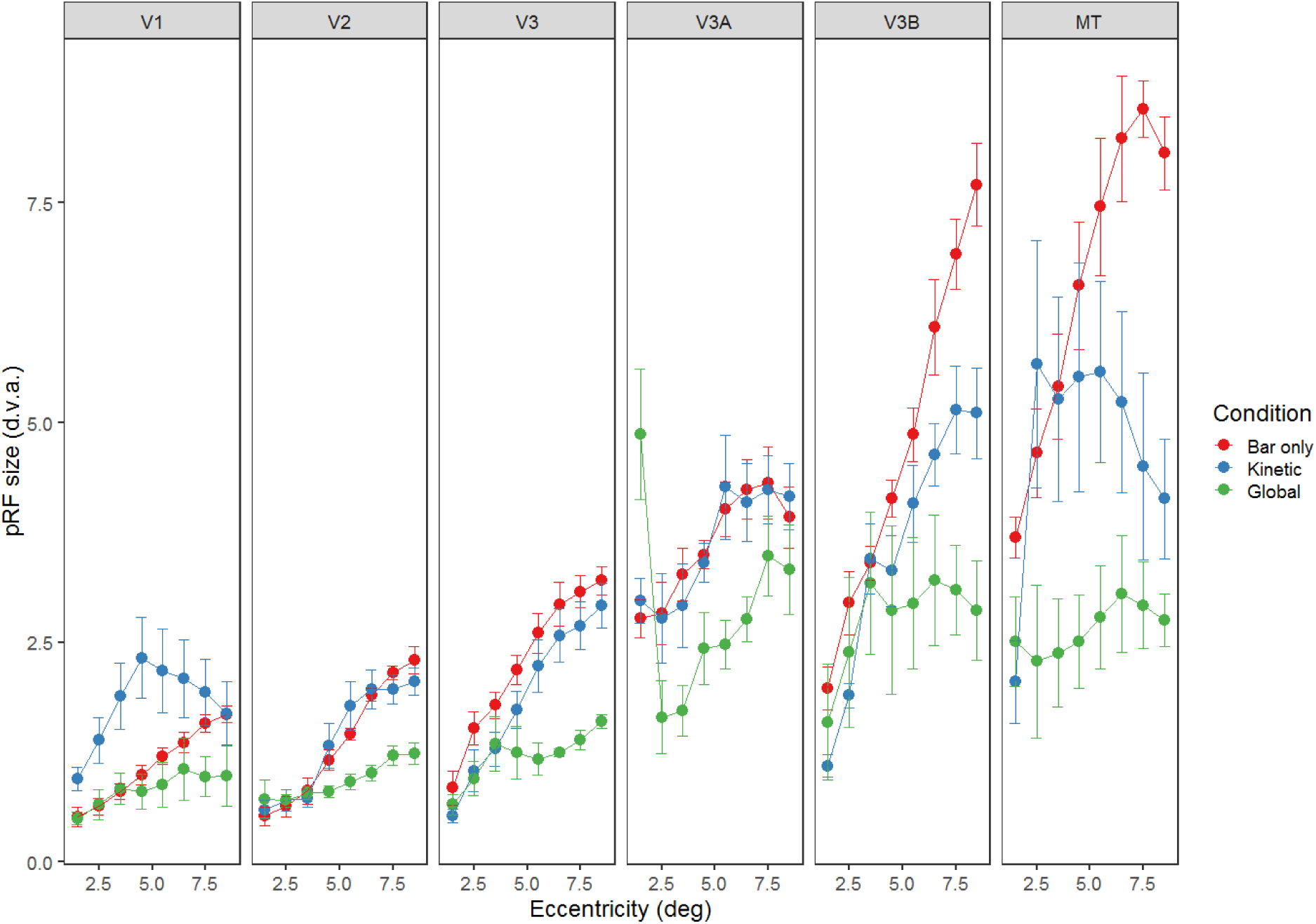
pRF size plotted against eccentricity for the different experimental conditions and brain regions in Experiment 1.

**Figure S3:**
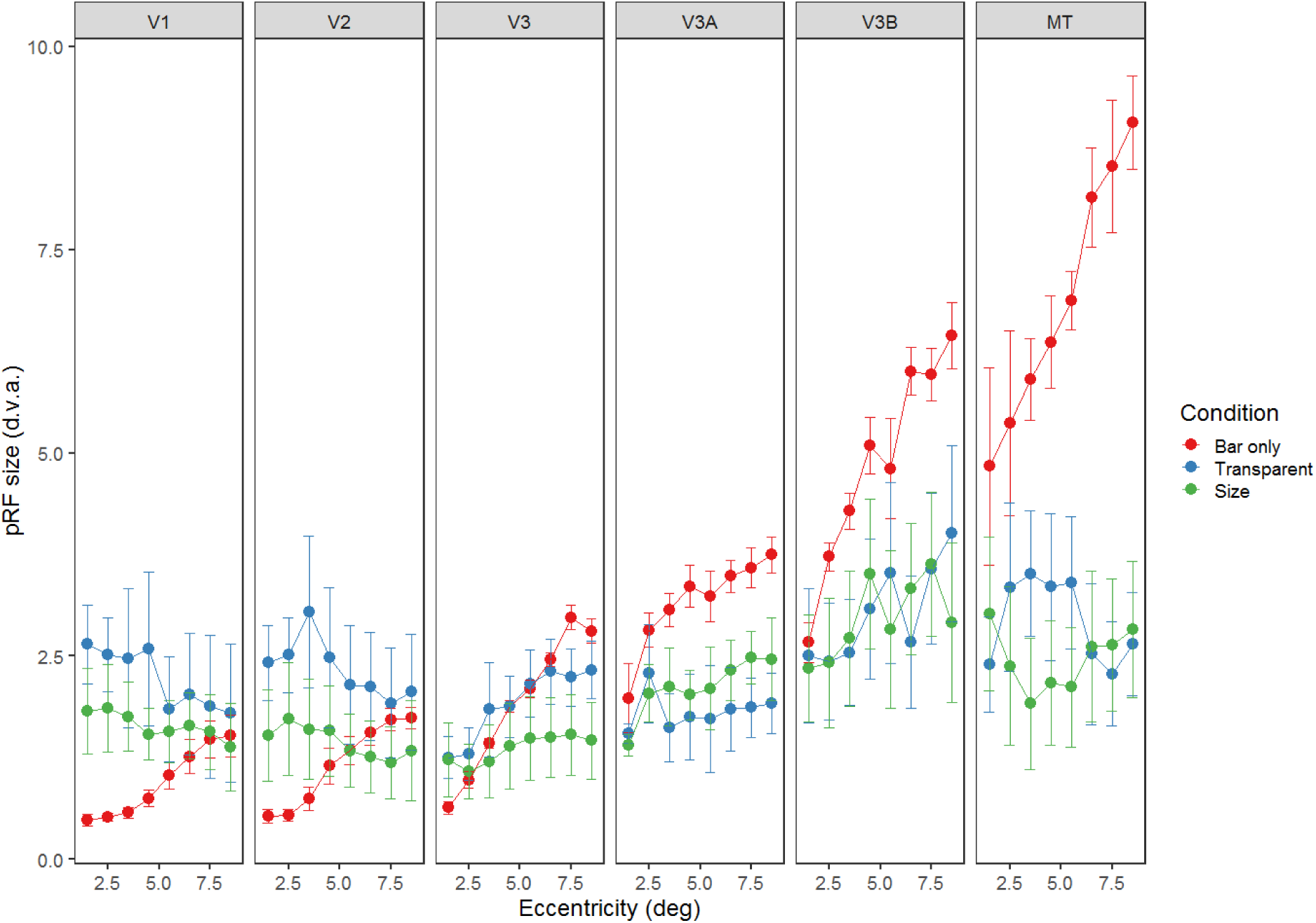
pRF size plotted against eccentricity for the different experimental conditions and brain regions in Experiment 2.

### Analyses using an R^2^ threshold of 0.1

As our experimental conditions often showed relatively weak and sparse responses, we used a fairly liberal R^2^ threshold of 0.05 in the main analyses. Here, we show the same analyses using a more conservative R^2^ threshold of 0.1. Overall, the results are highly comparable. We did not conduct formal statistical analysis due to the relatively high levels of missing data in some conditions and for some participants.

*Experiment 1*

**Figure S4.**
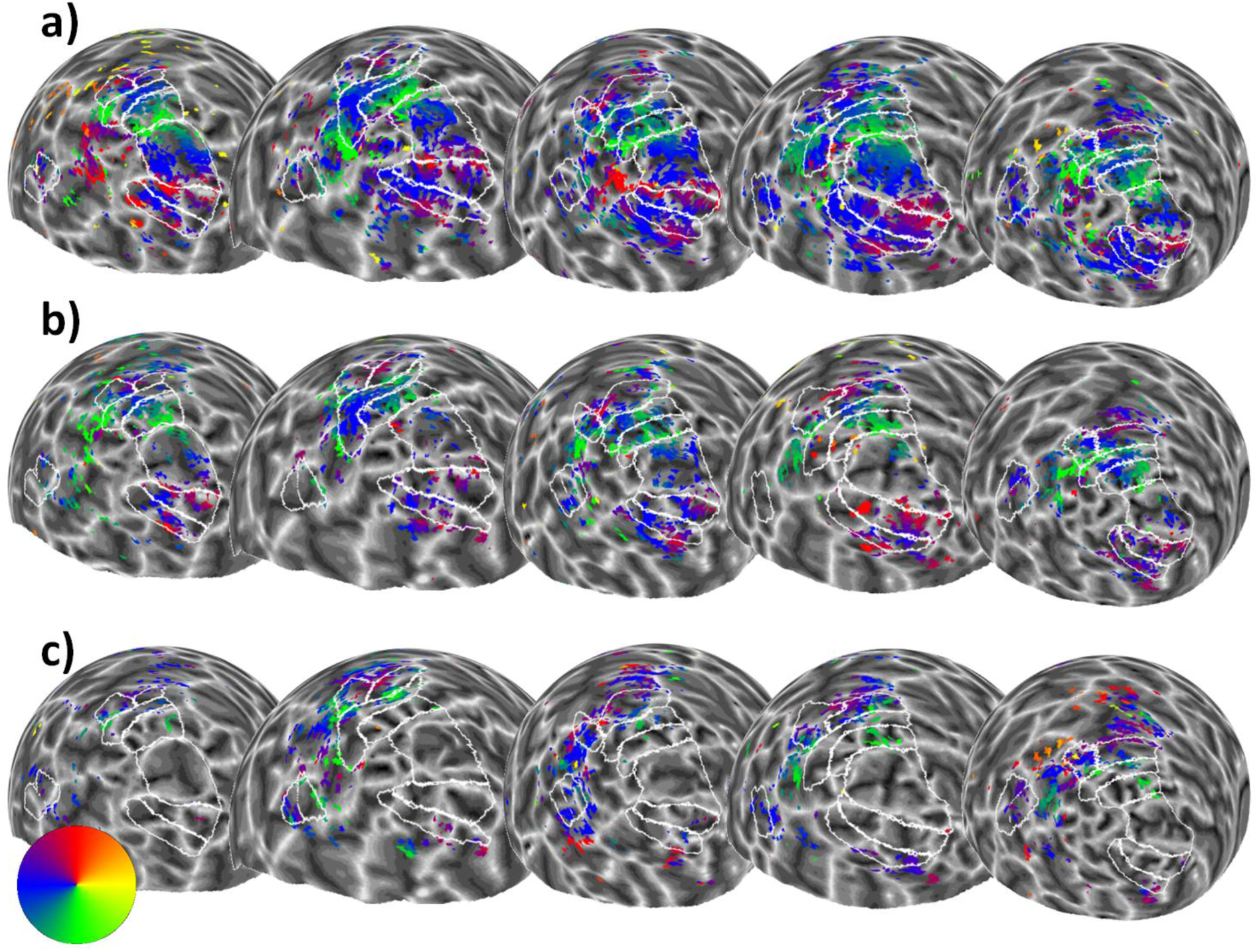
Sphere projection of polar angle data for the left hemispheres of all participants in Experiment 1 using an R^2^ threshold of 0.1. The colour of each vertex indicates the polar angle for the corresponding pRF centre (as indicated by the colour wheel). Each person’s data forms a column (subject 1 is on the far left, and subject 5 is on the far right), and stimulus condition forms a row. Manual delineations of visual areas V1, V2, V3, V3A, V3B and hMT+ (TO1/2) are shown. (**a**) Polar angle estimates for the ‘bar-only’ stimulus condition. (**b**) Polar angle estimates for the ‘kinetic’ stimulus condition. **(c)** Polar angle estimates for the ‘global’ stimulus condition.

**Figure S5.**
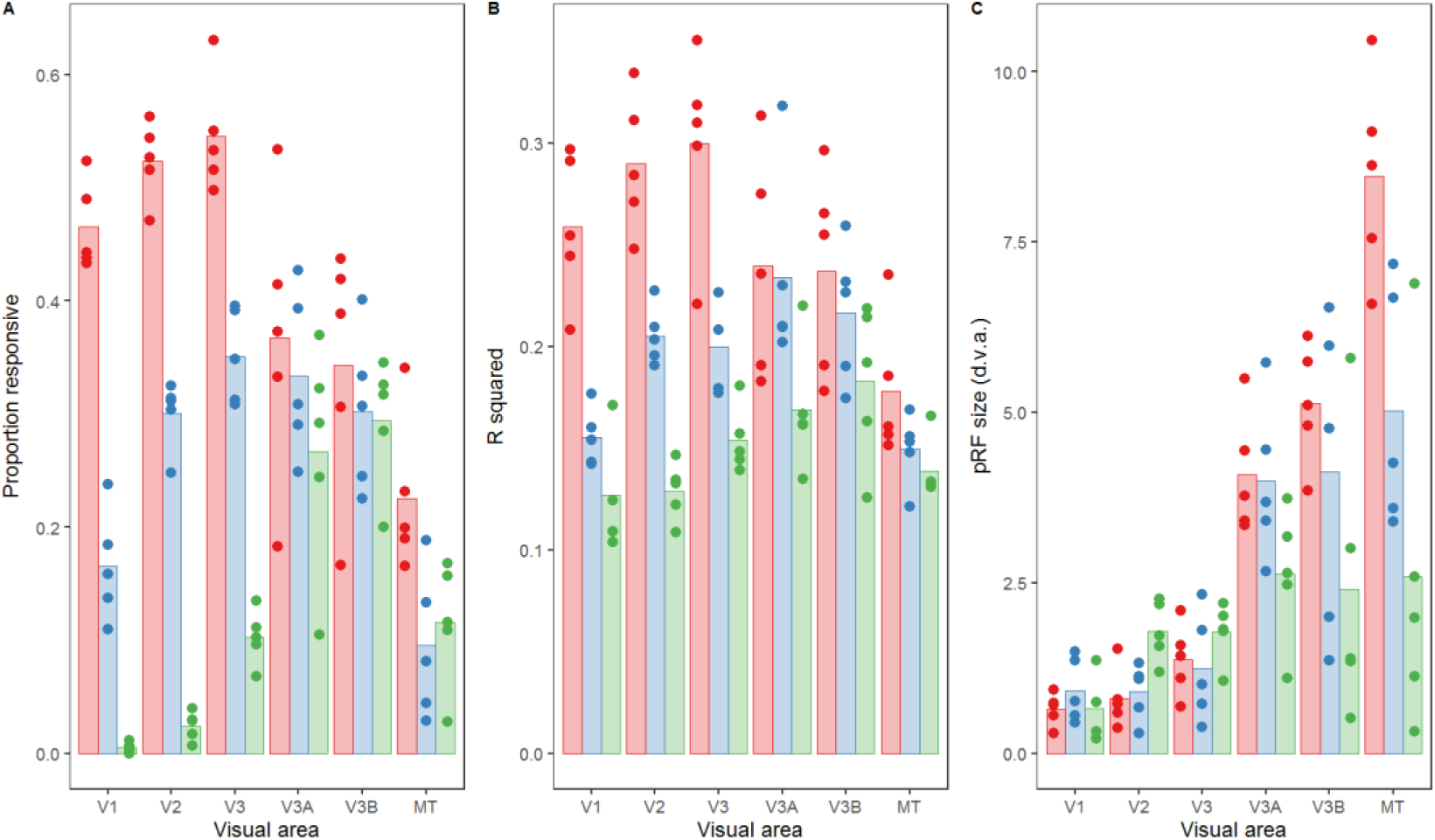
**(a)** Proportion of vertices responding, **(b)** goodness-of-fit and **(c)** pRF sizes for each condition and visual area in Experiment 1 with an R^2^ threshold of 0.1. The bars show the mean values across all subjects, and the points are individual data for each subject. Panel (a) plots the mean proportion of vertices responding for each subject, whereas (b) and (c) plot the median goodness-of-fit values and pRF sizes respectively. Subject 2 is missing data for the V1 global condition.

*Experiment 2*

**Figure S6.**
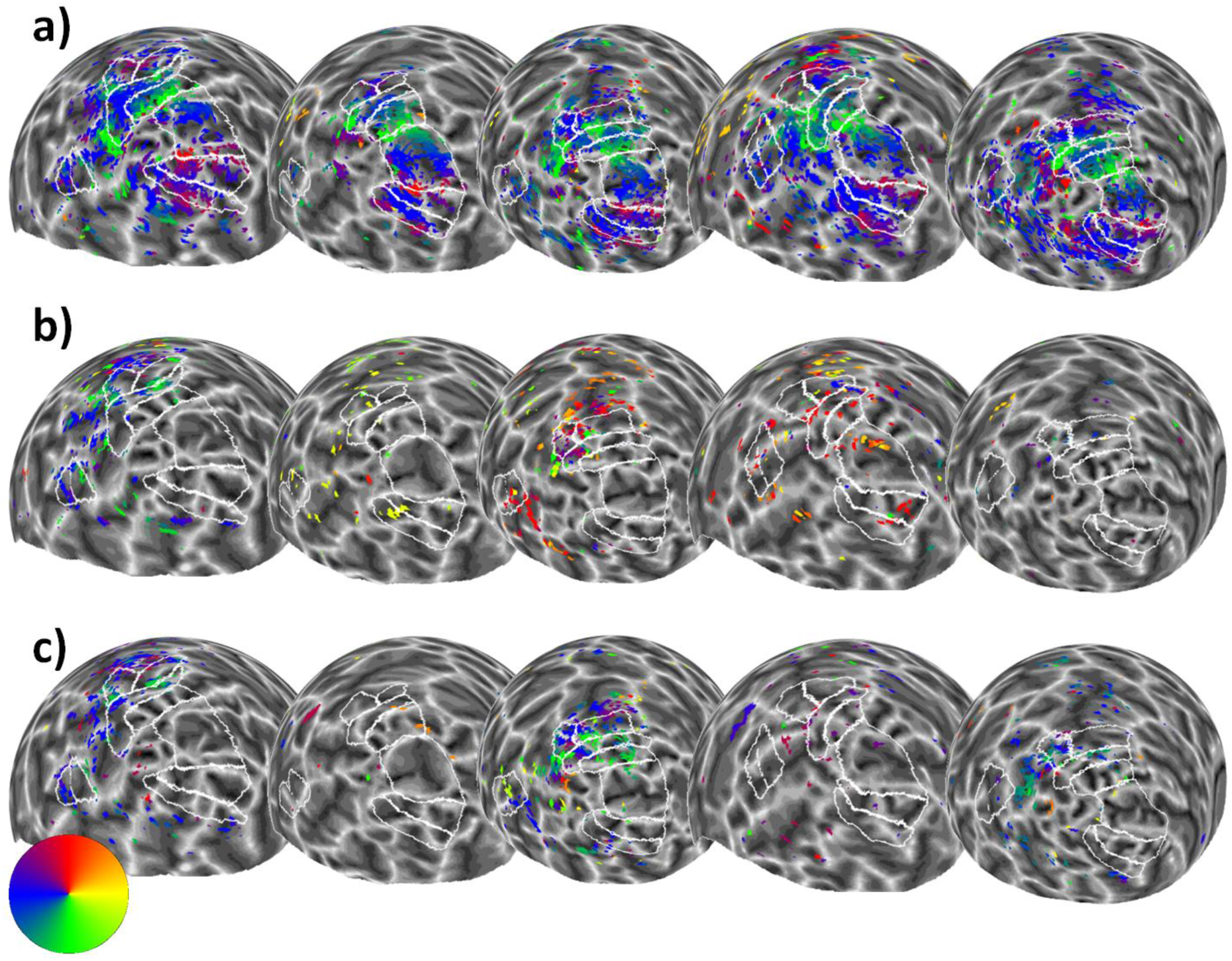
Sphere projection of polar angle data for the left hemispheres of all participants in Experiment 2 with an R^2^ threshold of 0.1. The colour of each vertex indicates the polar angle for the corresponding pRF centre (as indicated by the colour wheel). Each person’s data forms a column (subject 1 is on the far left and subject 5 is on the far right), and stimulus condition forms a row. Manual delineations of visual areas V1, V2, V3, V3A, V3B, and hMT+ (TO1/2) are shown (if the subject had taken part in Experiment 1, the delineations from this experiment were used). (**a**) Polar angle estimates for the ‘bar-only’ stimulus condition. (b) Polar angle estimates for the ‘transparent’ stimulus condition. (**c**) Polar angle estimates for the ‘size-defined’ stimulus condition.

**Figure S7.**
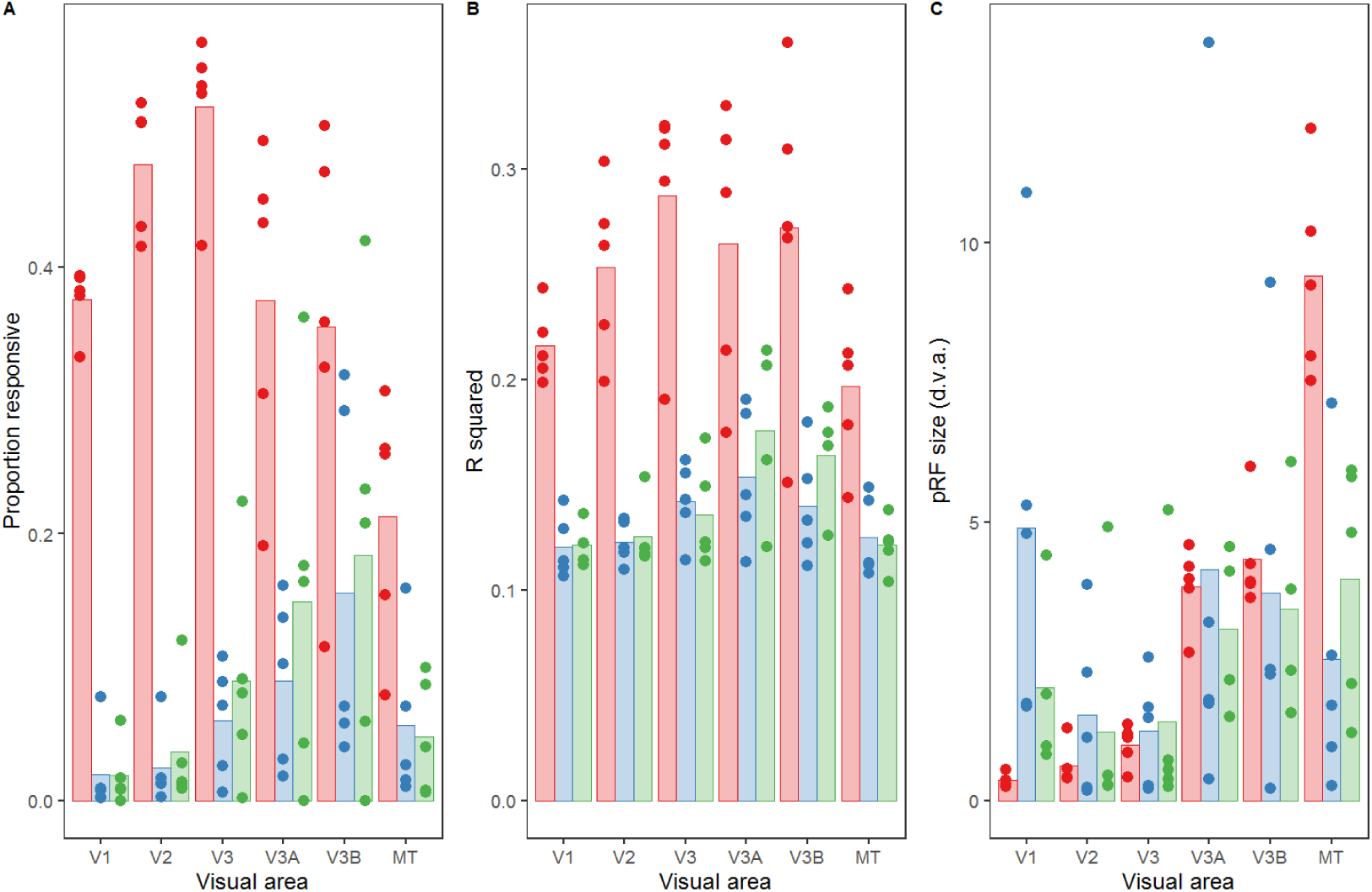
**(a)** Proportion of vertices responding, **(b)** goodness-of-fit and **(c)** pRF sizes for each condition (bar-only, transparent, and size-defined) and visual area in Experiment 2 with an R^2^ threshold of 0.1. The bars show the mean values across all subjects, and the points are individual data for each subject. In (a), this is the mean proportion of vertices responding for each subject, whereas for (b) and (c) these are the median goodness-of-fit values and pRF sizes respectively. Subject 2 is missing data for the size condition in V1, V3A and V3B.

